# Diverse lung challenges elicit a conserved monocyte-to-macrophage differentiation blueprint

**DOI:** 10.64898/2026.03.15.711867

**Authors:** Chrysante S. Iliakis, Wouter T’Jonck, Isobel C. Mouat, Sam Bankole, Jiawei Liang, Gareth-Rhys Jones, Justina Kulikauskaite, Matthew O. Burgess, Piotr Janas, Stefania Crotta, Simon L. Priestnall, Alejandro Suarez-Bonnet, Jürgen Schwarze, Andreas Wack, Calum C. Bain

## Abstract

Alveolar macrophages (AM) form a first line of defence to lung insults. These insults often lead to replacement of foetal-derived tissue-resident AMs (fAMs) by monocyte-derived AMs (mono-AMs) with different functionality, which impacts lung immunity long-term. However, whether these alterations are conserved or insult-specific is not well understood. Here, we show that respiratory syncytial virus (RSV) infection remodels the AM compartment long-term. We perform comparative analyses of fAMs and mono-AMs after RSV infection, influenza (IAV) infection and clodronate liposome administration to derive rules governing the functional and transcriptional changes in AM across insults. We identify a crucial tissue integration checkpoint of CD11b^+^ mono-AMs which involves acquisition of proliferative capacity and requires EGR2-mediated transcriptional rewiring. Mono-AMs elicited by RSV, influenza or clodronate liposomes have a hard-wired ontogeny-dependent stereotypical profile of transcription, metabolism and enhanced immunoreactivity despite a substantially different inflammatory environment during differentiation; accordingly, mono-AMs, whether elicited by infection or sterile depletion, protect from subsequent *S. pneumoniae* infection. By contrast, fAMs are more susceptible to subtle, context-dependent transcriptional alterations. Our findings highlight that long-term changes in the AM compartment show origin-dependent phenotypic divergence and that mono-AM integration follows a hard-wired trajectory fine-tuned by environmental factors.

## Introduction

Alveolar macrophages (AMs) are located in the broncho-alveolar space of the lung and under normal physiological conditions play a crucial role in lung homeostasis by regulating pulmonary surfactant, clearing dead and effete cells and maintaining the integrity of the epithelial barrier^1, 2^. They are also a critical component of the first line of defence against invading pathogens, as evidenced by increased susceptibility to infections and disease severity of mice and humans with AM deficiency^3, 4, 5, 6, 7, 8, 9^. In mice, AMs develop from foetal liver monocytes that colonize the lungs during alveologenesis and differentiate into AMs in response to paracrine GM-CSF and autocrine TGF-β^10, 11^. These cytokines converge to induce expression of the transcription factor PPAR-γ, which is indispensable for AM differentiation^12^. A number of other transcription factors have been implicated in AM differentiation, including CEBP/β^13^, EGR2^14, 15, 16^, Bhlhe40 and Bhlhe41^17^, Bach2^18^ and KLF4^19^.

Following viral and bacterial infection, AMs have been shown to adopt long-lasting transcriptional and functional changes^19, 20^. Because murine foetal-derived AMs (fAMs) are considered long-lived and require little replenishment from circulating monocytes under steady state conditions^21, 22, 23, 24, 25, 26, 27, 28^. early studies attributed these changes to tissue resident fAMs and considered this a form of innate imprinting or trained immunity^20, 21^. However, during lung infection, damage and/or inflammation, a common phenomenon is the highly coordinated and rapid depletion of fAMs^29, 30, 31^, sometimes referred to as the “macrophage disappearance reaction”^32^. Repopulation is essential to restore lung homeostasis and protective immune functions of AMs and can occur by proliferation of fAMs^27, 33, 34^ and/or recruitment of monocytes that differentiate locally into long-lived AM (mono-AMs)^14, 29, 30, 31, 35^; the relative contribution of these mechanisms can vary depending on the nature and magnitude of damage and inflammatory response. Our previous work and that of others has demonstrated that compelte AM repopulation following influenza A virus (IAV) infection is dependent on the recruitment of monocytes^29, 35^. Importantly, mono-AMs that arise following IAV infection have distinct transcriptional and functional properties compared with fAMs in the same lung, with differences persisting for months after viral clearance^29, 35^. Most notably, they have elevated responsiveness to Toll-like receptor (TLR) stimulation, which can be beneficial or deleterious upon subsequent infection depending on the context. For instance, the presence of IAV-elicited mono-AMs is protective against subsequent *Streptococcus pneumoniae* infection^29^. Similar observations were made for β-glucan-elicited mono-AMs, which are protective upon *Legionella pneumophila* challenge^36^. By contrast, the presence of IAV-elicited mono-AMs has been shown to cause enhanced disease severity during a subsequent IAV infection challenge ^35^. Hence, remodelling of the AM compartment following IAV infection has long-term consequences for lung immunity, but whether this reflects IAV-dependent environmental priming of mono-AMs or long-lasting, hard-wired features of mono-AMs, or both, remains unclear.

Infection with respiratory syncytial virus (RSV) is nearly universal in young children; over 80% are infected by the age of two, making RSV infection a common first lung insult in humans^37^. RSV is a single-stranded RNA virus that causes respiratory infections affecting upper and lower respiratory tracts, and is encountered repeatedly throughout life and in the majority of people causes only mild infection. However, severe RSV-induced bronchiolitis in young children can be associated with pulmonary sequelae later in life^38^. Although AMs are known to be a critical source of the early antiviral cytokine response during early phases of RSV infection in mouse models^39^, it remained largely unknown if and how a remodelling of the AM compartment, with potential long-term impacts, might occur after RSV infection.

Here, we used RSV infection in mice together with high-dimensional flow cytometric analysis, genetic lineage tracing and transcriptional profiling to show that RSV infection leads to marked and long-term remodelling of the AM compartment. We demonstrate that RSV induces cell death in fAM and that repopulation is a biphasic process; initially through *in situ* proliferation followed by replenishment by monocyte-derived cells. We show that mono-AMs progressively out-compete fAMs over the months following viral clearance and demonstrate that fAM and mono-AMs follow distinct transcriptional trajectories in the post infectious lung. Our transcriptional profiling reveals a checkpoint within mono-AMs governing their integration into the differentiated, self-renewing AM compartment, involving EGR2-mediated transcriptional rewiring and acquisition of a transient, suprahomeostatic proliferative state. We demonstrate that remodelling of AM composition alters functionality at population level, in large part due to the presence of mono-AMs which exhibit elevated cytokine and chemokine production, reduced mitochondrial and lipid metabolism and enhanced proliferation. Comparison with influenza A virus (IAV) infection and clodronate liposomes (clodr-lipo), a non-infectious method of eliciting mono-AMs, revealed that these functional differences were ‘hard-wired’, being conserved in mono-AMs across recruitment contexts. Finally, we demonstrate that clodronate-elicited mono-AMs are sufficient for protection upon subsequent challenge with *S. pneumoniae,* highlighting that the presence of AMs with monocyte origin confers prolonged functional changes to the AM compartment, even in the absence of prior lung infection.

## Results

### RSV infection induces long-term remodelling of the alveolar macrophage compartment

First, we assessed the infection dynamics in C57BL/6J mice upon intranasal inoculation with RSV (Human A2 strain). *Rsv-l* RNA levels (encoding the L protein) in whole lung tissue peaked at 2 days post-infection (dpi) and were undetectable by 14 dpi (**Extended data Fig. 1a**). RSV infection induced a robust type I interferon (IFN) response, indicated by increased expression levels of *Ifna1* and *Ifnb1* and IFN-stimulated genes (*Rsad2, Mx1, Pkr* and *Oas1a*) in whole lung tissue, as well as IFN-α and IFN-β in bronchoalveolar lavage fluid (BALF) (**Extended data Fig. 1b, c**). These responses were absent in mice inoculated with UV-inactivated RSV (UV-RSV) (**Extended data Fig. 1a-c**). RSV infection also resulted in an increased in the abundance of debris in the BALF, particularly at 7 days post-infection (**Extended data Fig. 1d**).

Next, we longitudinally characterised the myeloid cell landscape in the broncho-alveolar space. *Ms4a3*^Cre/+^*;Rosa26*^CAG-LSL-tdTomato/+^ mice, in which all cells deriving from granulocyte-monocyte progenitors (GMPs), including bone marrow monocytes and their progeny, can be traced, were analysed from 12 hours (hrs) to 2 months (mo) after RSV infection by flow cytometry. The myeloid compartment was selected by manual gating for CD3^−^CD19^−^NK1.1^−^(Lineage^−^) CD45^+^ cells expressing MerTK and/or CD11b, and initial analysis was based only on surface phenotype without making use of the lineage marker. Uniform Manifold Approximation and Projection (UMAP) analysis of concatenated data from all timepoints was performed to identify clusters (**Fig. 1a**). AMs were identified based on expression of SiglecF and CD11c, monocytes and monocyte-derived macrophages (MDMs) based on CD11b, Ly6C and MHCII expression, and neutrophils based on Ly6G expression (**Fig. 1a**). Over time, these subsets showed marked changes in abundance compared with control mice inoculated with UV-RSV (**Fig. 1b, c**). Neutrophil frequencies peaked at 1dpi, while monocytes and MDMs peaked later, reaching a maximum at 7 days post-infection (**Fig. 1b, c**). There were also dynamic changes in cell phenotype over time within the monocytes and MDM compartment; at 2 dpi, most cells expressed high levels of Ly6C, indicative of their recent derivation from blood Ly6C^hi^ monocytes (**Extended Data Fig. 1e, f**). At later timepoints, Ly6C expression was reduced and MHCII increased, indicative of monocyte-to-macrophage maturation (**Extended Data Fig. 1f**). Consistently, MerTK expression progressively increased within the Mono and MDM compartment over time (**Extended Data Fig. 1g**). AM numbers rapidly dropped within 12 hrs continuing to 2 dpi (**Fig. 1b, c**) paralleled by significantly reduced proliferation, assessed by Ki67 expression (**Fig. 1d**, upper panel), and increased cell death, assessed by Annexin V staining (**Fig. 1d**, lower panel). By 7 days post-infection, AM numbers, as well as the frequencies of proliferating and apoptotic cells, had returned to baseline (**Fig. 1b-d**). AMs also exhibited phenotypic changes during and after RSV infection. CD11b was transiently upregulated, peaking at 2 dpi (**Fig. 1e-f**), while MHCII expression increased and persisted as late as 2mo post-infection (**Fig. 1e-f**).

**Fig. 1.**
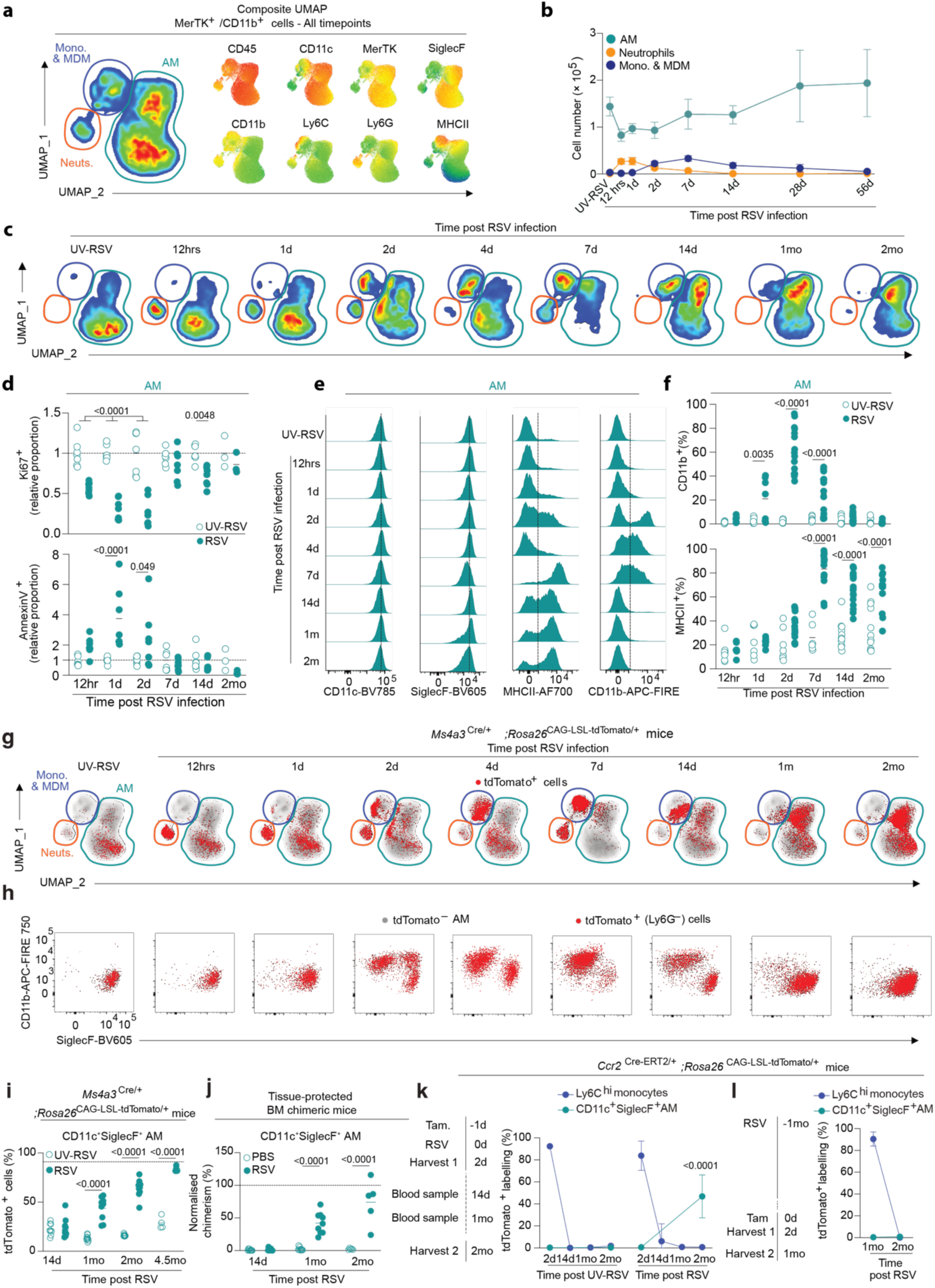
RSV infection induces long-term remodelling of the alveolar macrophage (AM) compartment characterized by engraftment and expansion of mono-Ams. **a.** Composite UMAP dimensionality reduction analysis of CD45^+^Lineage^−^ (CD3^−^NK1.1^−^CD19^−^) MerTK^+^ and/or CD11b^+^ cells within bronchoalveolar lavage fluid (BALF) from *Ms4a3*^Cre/+^;*Rosa26*^LSL-CAG-tdTomato/+^ mice harvested at 12hrs, 1, 2, 4, 7, 14d, 1mo and 2mo post-RSV infection or from UV-RSV controls. Colours within the contour plot (left) reflect cell abundance, while colours in feature dot plots (right) reflect relative expression of the indicated markers. UMAPs were generated using concatenated data from *n* = 3 mice per time point. **b.** Absolute numbers of indicated cell subsets in BALF at the indicated time points. Symbols represent average count and error bars represent ± 1SEM. Data pooled from two independent experiments with *n* = 3-7 mice per group; UV-RSV group pooled from multiple time points. **c.** UMAP dimensionality reduction analysis from **a** (left) segregated for each timepoint post-infection. **d.** Proportion of Ki67^+^ (top panel) or Annexin V^+^ (bottom panel) CD11c^+^SiglecF^+^ BALF AMs at indicated time points after RSV or UV-RSV inoculation. Data presented as relative proportion to time-matched UV-RSV group. Symbols represent individual mice. Data pooled from two independent experiments with *n* = 3-7 mice per group. Statistical significance calculated by two-way ANOVA. **e.** Representative flow cytometry histograms of CD11c, SiglecF, MHCII and CD11b expression by CD11c^+^SiglecF^+^ BALF AM at the indicated time post-RSV infection. **f.** Proportion of CD11b^+^ (top panel) or MHCII^+^ (bottom panel) CD11c^+^SiglecF^+^ BALF AMs at indicated time points after RSV or UV-RSV inoculation. Data presented as relative proportion to time-matched UV-RSV group. Points represent individual mice. Data pooled from two to six independent experiments with *n* = 7-24 mice per group. Statistical significance calculated by two-way ANOVA. **g.** Expression of tdTomato by MerTK^+^ and/or CD11b^+^ (CD45^+^CD3^−^NK1.1^−^CD19^−^) cells from BALF of *Ms4a3*^Cre/+^;*Rosa26*^LSL/CAG-tdTomato/+^ mice from **a**. Grey contour plots represent the entire MerTK^+^ and/or CD11b^+^ compartment pooled from all timepoints with red dots representing tdTomato^+^ cells present only at the indicated time point. **h.** Representative flow cytometry plots showing expression of CD11b and SiglecF by tdTomato^−^ AM and Ly6G^−^tdTomato^+^ cells within the MerTK^+^ and/or CD11b^+^ compartment within BALF. **i.** Proportion of tdTomato^+^ cells amongst CD11c^+^SiglecF^+^ AMs in BALF at indicated time points after RSV or UV-RSV inoculation. Symbols represent individual mice. Data pooled from two independent experiments with *n* = 4-9 mice per group. Dotted line represents average proportion across all timepoints of labelled circulating monocytes in blood. Statistical significance calculated by two-way ANOVA. **j.** Chimerism of CD11c^+^SiglecF^+^ AMs within BALF from tissue-protected bone marrow chimeric mice normalised to blood Ly6C^hi^ monocyte chimerism at the indicated time points post infection. Symbols represent individual mice. Dashed line indicates average chimerism in blood monocytes. Data pooled from two independent experiments with *n* = 5-9 mice per group. Statistical significance by two-way ANOVA. **k.** Proportion of tdTomato^+^ Ly6C^hi^ monocytes in blood and CD11c^+^SiglecF^+^ AM in BALF obtained from *Ccr2*^Cre-ERT2*/+*^*;Rosa26*^CAG-LSL-tdTomato/+^ mice given a single dose of tamoxifen 24 hrs prior to inoculation with UV-RSV or RSV. Blood monocytes were assessed at all indicated timepoints. AMs in BALF were assessed at 2 days and 2 months post-infection. Symbols represent mean ± S.D.. Data pooled from one (UV-RSV) or two independent (RSV) experiments with *n* = 2-7 mice total per group. Statistical significance calculated by two-way ANOVA. **l.** Proportion of tdTomato^+^ Ly6C^hi^ monocytes in blood and CD11c^+^SiglecF^+^ AM in BALF from *Ccr2*^Cre-ERT2*/+*^*;Rosa26*^CAG-LSL-tdTomato/+^ mice inoculated with UV-RSV or RSV and administered tamoxifen one month later. Blood monocytes and AMs were assessed two days and one month after administration of tamoxifen (two months post-infection) Data shown pooled from two independent experiments with *n* = 7-9 mice in total per group.

We leveraged the lineage tracing of bone marrow monocytes in *Ms4a3*^Cre/+^*;Rosa26*^CAG-LSL-tdTomato/+^ mice to determine definitively the trajectory of monocytes over time after RSV infection. Lack of tdTomato positivity in this context can be interpreted as derivation from *Ms4a3*-independent progenitors, including foetal liver monocytes. In *Ms4a3*^Cre/+^*;Rosa26*^CAG-LSL-tdTomato/+^ mice administered UV-RSV, the majority of AMs was not labelled, in keeping with their predominantly foetal origin (**Fig. 1g, h**). As expected, tdTomato^+^ cells were abundant within neutrophil, monocyte and MDM compartments between 12 hrs and 7dpi (**Fig. 1g**). However, by 14dpi, the positioning of tdTomato^+^ cells was on the boundary between monocytes/MDMs and the AM compartment (**Fig. 1g**, 7dpi); indeed, within Ly6G^−^ tdTomato^+^ cells, a continuum was observed between 14d and 1mo from CD11b^+^ to SiglecF^+^ and CD11c^+^ compartments (**Fig. 1h** and **Extended Data Fig. 1h**), indicative of differentiation of monocytes to monocyte-derived AMs (mono-AMs). By 1 mo post infection, the vast majority of tdTomato^+^ cells were found in the SiglecF^+^ AM gate. Indeed, we found a progressive increase in tdTomato^+^ cells amongst SiglecF^+^ cells after 14 dpi (**Fig. 1i**). Notably, by 4.5 mo post-infection tdTomato labelling within the AM compartment reached similar levels to that seen in Ly6C^hi^ monocytes in blood; a phenomenon not seen in recipients of UV-RSV or aged unmanipulated *Ms4a3*^Cre/+^*;Rosa26*^CAG-LSL-tdTomato/+^ mice (**Fig. 1i** and **Extended Data Fig. 1i**).These data indicate that the initial repopulation following RSV-induced AM loss occurs largely independently of monocytes, but that monocyte-derived AMs arise after viral clearance and ultimately come to dominate the alveolar niche. Consistent with other models^7, 21, 30, 31, 40^, we found that mono-AMs expressed lower levels of SiglecF which had not reached levels of foetal-derived alveolar macrophages (fAM) by 2 mo post infection (**Extended Data Fig. 1j**). Importantly, the contribution of monocyte-derived cells was not entirely responsible for the MHCII expression by the AM compartment in the post-RSV lung, as both tdTomato^+^ mono-AM and tdTomato^−^ fAMs displayed high levels of MHCII (**Extended Data Fig. 1j**), suggesting this phenotypic feature is driven by the post-RSV environment. Finally, to validate and complement these fate mapping studies, we assessed replenishment kinetics in tissue protected bone chimeric mice in which wild type CD45.1^+^/CD45.2^+^ mice were irradiated with their lungs (and upper body) protected with lead before being reconstituted with congenic CD45.2^+^ bone marrow. At least eight weeks later, mice were infected or not with RSV and the chimerism of AMs assessed longitudinally. Consistent with our results in *Ms4a3*^Cre/+^*;Rosa26*^CAG-LSL-tdTomato/+^ mice, we detected progressive accumulation of bone marrow mono-AM following RSV but not in control mice (**Fig. 1j**).

The accumulation of mono-AM following RSV infection could be explained by continual recruitment of monocytes after infection, or could reflect the recruitment of monocytes during infection that differentiate into mono-AM and subsequently outcompete fAM counterparts. While useful for tracing replenishment dynamics, the constitutive nature of labelling in *Ms4a3*^Cre/+^*;Rosa26*^CAG-LSL-tdTomato/+^ mice cannot distinguish between these alternatives. Thus, we turned to an inducible system in which CCR2^+^ cells, including Ly6C^hi^ monocytes, can be temporally labelled through administration of tamoxifen. Delivery of a single dose of tamoxifen to *Ccr2*^Cre-ERT2*/+*^*;Rosa26*^CAG-LSL-tdTomato/+^ mice leads to efficient and irreversible labelling of the Ly6C^hi^ monocyte population which decays over time, consistent with their short half-life and hence replacement with non-labelled cells (**Extended Data Fig. 1k**). Importantly, there was no discernible labelling of AM early post infection in these same mice (**Extended Data Fig. l**), demonstrating the utility of this system for ‘time stamping’ monocytes. Thus, we used this system to label Ly6C^hi^ monocytes in blood one day prior to RSV infection and assessed the presence of tdTomato^+^ AMs 2d (3 days post tamoxifen) and at 2mo post-infection (**Fig. 1k**). Consistent with our longitudinal profiling in *Ms4a3*^Cre/+^*;Rosa26*^CAG-LSL-tdTomato/+^ mice, labelling CCR2^+^ cells prior to infection led to the presence of tdTomato^+^ Ly6C^hi^ monocytes within the BALF at 2 dpi (**Fig. 1k**). Importantly, even in the context of acute RSV, no AMs were labelled by tamoxifen administration (**Fig. 1k** & **Extended Data Fig. 1m**). When the BALF was examined at 2mo post-RSV, we found that ∼60% of AMs were now tdTomato^+^ and little, if any, Ly6C^hi^ monocytes could be found in the BALF (**Fig. 1k, Extended Data Fig. 1n**). Again, tdTomato^+^ AM displayed lower levels of SiglecF expression compared with their tdTomato^−^counterparts (**Extended Data Fig. 1o**). To rule out continual replacement by recruited monocytes, we next assessed AM labelling when monocytes were pulse-labelled at 35 dpi, the point at which a monocyte-derived cell engraftment was seen in the AM compartment but before complete outcompetition as observed in BM chimeric and *Ms4a3*-based systems (**Fig. 1i, j**). However, while blood monocytes were labelled in this experiment, when assessed at 2mo post-infection there were essentially no tdTomato^+^ AMs (**Fig. 1l**). Taken together, these data support the idea that monocytes elicited during acute RSV infection differentiate progressively from a CD11b^+^ stage into *bona fide* CD11b^−^ SiglecF-expressing AMs that come to outcompete their foetally-derived counterparts. We refer to this process of phenotypic convergence and acquisition of self-renewal capacity as “integration”.

### Acquisition of supra-homeostatic proliferative capacity is a crucial tissue integration checkpoint

Having established the kinetics of mono-AM emergence, we next sought to understand the transcriptional changes that occur during this process. Thus, we examined the transcriptional profile of CD11b- or SiglecF-defined populations representing distinct stages of the MDM to mono-AM trajectory over time using bulk RNA-sequencing (CD11b^+^, SiglecF^lo^ and SiglecF^hi^ cells from 14d, 1 and to 2mo post-infection) (**Fig. 2a**). Consistent with progressive integration over time, there was a gradual downregulation of monocyte-associated genes such as *Ccr2* and *Cd93,* whereas *Siglecf* and genes associated with an AM signature, including *Pparg, Marco, Fabp1* and *Epcam* were found to be gradually upregulated (**Fig. 2b**). In line with previous results (**Fig. 1e-f**), MHCII-associated genes remained elevated in virus-experienced SiglecF^hi^ cells isolated from RSV-infected mice compared to UV-RSV controls. To validate our *a priori* division on CD11b/SiglecF, we derived transcriptional signatures for each of the “integration stages” (CD11b^+^, SiglecF^lo^, and SiglecF^hi^) and performed module scoring of these within a publicly available single cell RNA-seq (scRNA-seq) dataset of monocytes/macrophages in the context of influenza A virus (IAV) infection^36^. In that dataset, monocytes, *bona fide* AM, and a subset of ‘transitional macrophages’ were defined, and trajectory inference analysis suggested a developmental trajectory from monocytes to AM through the transitional macrophage subset^36^. Consistent with these findings, we observed that the CD11b^+^ AM signature from our RSV dataset at 14 dpi directly overlayed onto ‘transitional macrophage subsets’ in the IAV dataset, while the signatures of SiglecF^lo^ mono-AMs at 1mo and SiglecF^hi^ at 2mo cells after RSV infection showed enrichment within AM subsets, although occupying somewhat discrete areas of the UMAP, consistent with them remaining transcriptionally distinct (**Fig. 2c**). Pathway analysis suggested large metabolic shifts across the CD11b^+^ > SiglecF^lo^ > SiglecF^hi^ trajectory; a finding we confirmed using complementary flow cytometry-based metabolic analyses, including glucose and lipid uptake, lipid context and peroxisome and mitochondrial mass (**Fig. 2d, Extended Data Fig. 2a**). In particular, CD11b^+^ cells displayed a metabolic profile that was highly distinct from cells expressing SiglecF. Of note, the largest number of DEGs was found at the transition from CD11b^+^ AM to SiglecF^lo^ AM (**Fig. 2e**). Of these, many were genes associated with cell cycle, which was reflected in the pathways enriched by overrepresentation analysis of upregulated DEGs (**Fig. 2f**). This suggested that part of the macrophage integration process involves entry into a proliferative state. To validate this, we determined proliferative rates by assessing expression of Ki67 at 1mo after RSV infection across monocyte/macrophage subsets defined by CD11b and SiglecF expression. We found that SiglecF^hi^ AM at 1mo after RSV infection had equivalent levels of proliferation to their counterparts in control (UV-RSV) mice (**Fig. 2g, Extended Data Fig. 2b**). However, a greater proportion of SiglecF^lo^ mono-AM expressed Ki67, indicating enhanced proliferation within this compartment. By contrast, only a very small proportion of CD11b^+^ mono-AM that had not yet adopted SiglecF expression were Ki67^+^, indicating that an elevated proliferation rate was not a global feature of monocyte-derived cells (**Fig. 2g**). Equivalent findings were obtained when using a short-term (2h) EdU pulse-chase approach (**Fig. 2h, Extended Data Fig. 2c**). Taken together, these data support the premise that differentiation from a SiglecF^−^CD11b^+^ state to a SiglecF^hi^ state involves acquisition of a highly proliferative SiglecF^lo^ state.

**Fig. 2.**
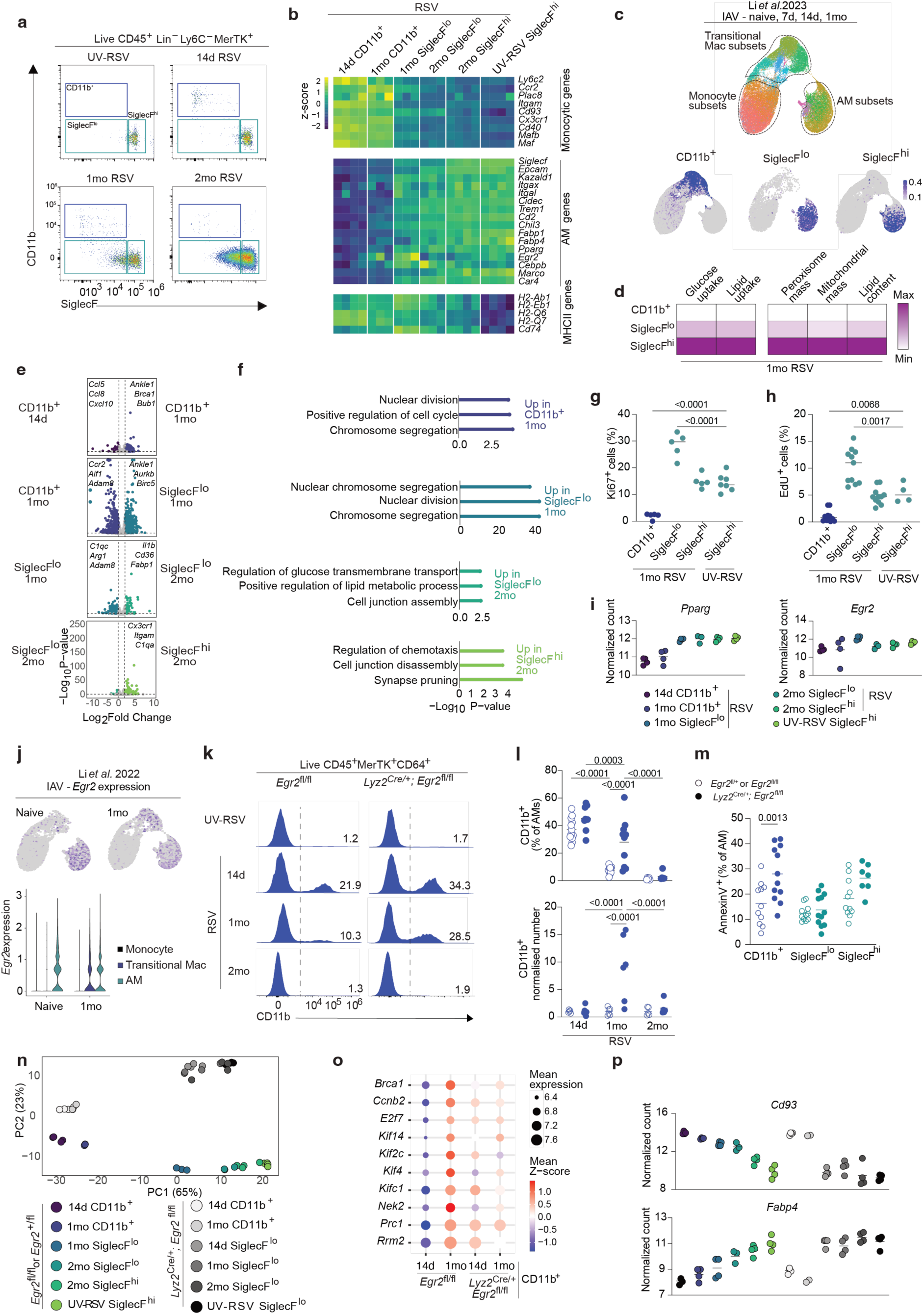
Acquisition of supra-homeostatic proliferative capacity is a crucial tissue integration checkpoint. **a.** Gating strategy used for sorting of identify AM subpopulations after RSV infection. Mono-AMs were identified as SiglecF^lo^ and stratified according to CD11b expression, while fAMs were defined as SiglecF^hi^ CD11b^−^. Cells were gated upstream as MerTK^+^CD64^+^CD11c^+^. **b.** Relative expression level (z-score) of selected monocyte- and AM-, and MHCII-associated genes from AM subpopulations at different integration stages in *Egr2*^flox/flox^ control mice inoculated with RSV or UV-RSV. UV-RSV samples pooled from 14d and 1mo timepoints. Each column represents one biological replicate. Subpopulations arising from equivalent timepoints were isolated from the same mouse. **c.** UMAP of scRNA-seq data (from Li *et al.* 2023, Science Immunology^35^) of monocytes and macrophages isolated from naïve or IAV-infected mice harvested at 7d, 14d or 1mo post-infection. Landmark gene signatures were used to assign clusters to three superclusters: monocytes, transitional macrophages and AM. Grey-to-blue colour intensity represents the average log_2_ fold change of subset-defining gene signatures from CD11b^+^, CD11c^+^CD11b^−^SiglecF^lo^ and CD11c^+^CD11b^−^SiglecF^hi^ AM subsets in *Egr2*^flox/flox^ control mice. **d.** Relative peroxisome mass (Pex14-CL594), Mitochondrial mass (mitotracker orange), intracellular neutral lipid content (BODIPY493/503), long-chain fatty acid uptake (BODIPY C-16) and glucose uptake (2-NBDG) in CD11b^+^, SiglecF^lo^ and SiglecF^hi^ AM subsets at 1mo post-RSV. Values normalized within one analyte across subpopulations. **e.** RNAseq-based pairwise gene expression comparison of sorted AM subsets defined by CD11b and SiglecF expression as indicated in (a), isolated from RSV-infected *Egr2*^flox/flox^ mice 14d, 1mo, and 2mo post-infection or inoculated with UV-RSV. **f.** Over-representation analysis of genes upregulated in integrating AM subsets delineated in **e**, determined by genes significantly upregulated within each pairwise comparison. **g.** Frequency of Ki67^+^ cells within CD11b-/SiglecF-defined AM subsets isolated from mice 1mo post-inoculation with RSV or UV-RSV. Symbols represent biological replicates. Data show from one representative experiment of two with *n=*5-7 mice per group. Statistical significance calculated by one-way ANOVA. **h.** Frequency of EdU^+^ cells within CD11b-/SiglecF-defined AM subsets isolated from mice 1mo post-inoculation with RSV or UV-RSV. EdU administered 2hrs before harvest. Symbols represent biological replicates. Data pooled from two independent experiments with total *n* = 4-11 mice per group. Statistical significance calculated by one-way ANOVA. **i.** Expression level of *Pparg* and *Egr2* in *Egr2*^flox/flox^ controls across CD11b/SiglecF-defined AM subsets at the indicated timepoints. **j.** Expression level of *Egr2* within monocytes, transitional macrophages and AM clusters described in **c** from Li *et al.* 2023, Science Immunology^35^ in naive or IAV-infected mice 1mo post-infection. **k.** Representative histograms of CD11b expression within live CD45^+^Lin^−^Ly6C^−^MerTK^+^ cells obtained from RSV-infected *Lyz2*^Cre/+^;*Egr2*^flox/flox^ mice or *Egr2*^flox/flox^ littermates controls infected at 14d, 1mo and 2mo post-infection. **l.** Frequency (*top*) and normalised number (*bottom*) plots of BALF CD11c^+^CD11b^+^ AMs. Symbols represent individual mice. Total of *n* = 5-7 mice per group, data shown pooled from two independent experiments. Statisical significance calculated using two-way ANOVA. **m.** Proportion of Annexin V^+^ CD11b- and SiglecF-defined AM subsets isolated from *Egr2*^flox/flox^ and *Lyz2*^Cre/+^;*Egr2*^flox/flox^ mice at 1mo after RSV infection. Data pooled from two independent experiments with *n* = 11 per group. Symbols represent individual mice. Statistical significance calculated using two-way ANOVA. **n.** PCA plot of RNA-seq data of CD11b- and SiglecF-defined AM subsets isolated from BALF of *Egr2*^flox/flox^ control and *Lyz2*^Cre/+^;*Egr2*^flox/flox^ mice inoculated with RSV or UV-RSV and harvested at the indicated timepoints. **o.** Dot plot displaying the mean expression level and mean z-score of selected cell cycle-associated genes from from *Egr2*^flox/flox^ and *Lyz2*^Cre/+^;*Egr2*^flox/flox^ at 14d and 1mo post-RSV **p.** Expression level of *Cd93* and *Fabp4* in integrating AM subsets from RNA-seq data in (n) from *Egr2*^flox/flox^ and *Lyz2*^Cre/+^;*Egr2*^flox/flox^ mice.

To understand the putative molecular regulators of this differentiation, we assessed expression of known transcription factors controlling AM differentiation, including PPARγ and EGR2, within our RNA-seq dataset. Whereas PPARγ expression increased progressively across the CD11b^+^ > SiglecF^hi^ trajectory, EGR2 showed a distinct expression pattern with a spike of expression which coincided with the transition into a highly proliferative state (i.e. within CD11b^−^ SiglecF^lo^ mono-AM) (**Fig. 2i**). Interestingly, whereas EGR2 was a specific feature of mature AM in naïve mice within the Li *et al*. scRNA-seq dataset, expression appeared in the ‘transitional macrophages’ in the context of IAV (**Fig. 2j**)^35^. Our previous work showed that EGR2 controlled AM repopulation in the context of sterile injury using the bleomycin-induced injury and fibrosis model^14^, however if and how EGR2 regulates monocyte-to-AM integration, particularly during viral infection, remained unexplored. Therefore, we examined if EGR2 deficiency altered mono-AM integration by infecting *Lyz2*^Cre/+^;*Egr2*^fl/fl^ mice (which lack *Egr2* within the myeloid compartment) and their *Lyz2*^+/+^;*Egr2*^fl/fl^ littermate controls (henceforth referred to as *Egr2*^fl/fl^ controls) with RSV and profiled BALF cells at 14d, 1mo, and 2mo post-infection. Of note, due to the dependence of SiglecF expression on EGR2^14, 16^ in these experiments mature AM could only be identified as CD11b^−^, making distinction of mono-AM from fAM more complex. Following RSV infection, the numbers of CD11b^−^ AM were comparable between both genotypes (**Extended Data Fig. 2d**), indicating repopulation by *in situ* proliferation is unaffected by *Egr2* deficiency. Accumulation of monocyte-derived CD11b^+^ macrophages at 14d was equivalent between *Lyz2*^Cre/+^;*Egr2*^fl/fl^ mice and their *Egr2*^fl/fl^ littermate controls (**Fig. 2k, l**). However, whereas at 1mo post-infection the CD11b^+^ macrophage population size had reduced to homeostatic levels in the *Egr2*^fl/fl^ controls, there was a marked persistence of these cells in *Lyz2*^Cre/+^;*Egr2*^fl/fl^ mice (**Fig. 2k, l**). Assessment of EGR2 expression by flow cytometry confirmed the loss of EGR2 expression in these persisting cells (**Extended Data Fig. 2e**). By 2mo post-infection the CD11b^+^ population had returned to baseline levels in both *Egr2*^fl/fl^ controls and *Lyz2*^Cre/+^;*Egr2*^fl/fl^ mice (**Fig. 2k, l**)

This raised the possibility that the mono-AM integration process is either defective, ultimately resulting in loss of the cells, or delayed, in the absence of EGR2 in the context of a viral infection. To distinguish between these alternatives, we first assessed the frequency of apoptotic cells across AM compartments in *Egr2*^fl/fl^ and *Lyz2*^Cre/+^;*Egr2*^fl/fl^ mice by Annexin V staining. CD11b^+^ cells, but not SiglecF^lo^ or SiglecF^hi^ cells, showed significantly elevated frequencies of Annexin V^+^ cells in *Lyz2*^Cre/+^;*Egr2*^fl/fl^ mice compared with *Egr2*^fl/fl^ controls (**Fig. 2m**), suggesting these cells have a defective capacity to survive in the lung long-term. As resident foetal-derived cells in *Lyz2*^Cre/+^;*Egr2*^fl/fl^ mice lack SiglecF expression^14, 16^, progression to a SiglecF^lo^ state could not be used to determine CD11b^+^ cell differentiation capacity by flow cytometry. We therefore examined the ‘stages of integration’ of mono-AMs over time by bulk RNA-seq in *Lyz2*^Cre/+^;*Egr2*^fl/fl^ mice and *Egr2*^fl/fl^ controls (see **Extended Data Fig. 2f** for sorting strategy). Of note, SiglecF^hi^ cells in *Lyz2*^Cre/+^;*Egr2*^fl/fl^ mice, representing Cre escapees, were not sorted for analysis. We observed that even the earliest stages of differentiation were changed in *Lyz2*^Cre/+^;*Egr2*^fl/fl^ mice, with CD11b^+^ AM remaining transcriptionally similar between 14d and 1mo compared with controls, as assessed by their relative transcriptional positioning by PCA (**Fig. 2n**). This contrasted with *Egr2*^fl/fl^ controls, where CD11b^+^ cells shifted towards SiglecF^hi^ resident AMs transcriptionally along principal component 1 (**Fig 2n**). Of the 327 genes differentially expressed in CD11b^+^ cells from *Egr2*^fl/fl^ controls between 14d and 1mo post-infection, only 40% were also differentially expressed in *Lyz2*^Cre/+^;*Egr2*^fl/fl^ mice (**Extended Data Fig. 2g**). As pathways associated with cell cycle were of the most upregulated at this stage in *Egr2* sufficient cells, we focussed on genes associated with cell cycle. Despite the low overall levels of proliferation in the CD11b^+^ compartment, the expression levels of several proliferation genes were increased robustly between 14d and 1mo post-infection in *Egr2*^fl/fl^ mice (**Fig. 2o**). However, this same pattern was not seen in *Lyz2*^Cre/+^;*Egr2*^fl/fl^ mice (**Fig. 2o**). Furthermore, the proliferative rate in CD11b^−^ AM from *Lyz2*^Cre/+^;*Egr2*^fl/fl^ mice was equivalent compared with UV-RSV controls, indicating that highly proliferative monocyte-derived cells do not contribute to this compartment (**Extended Data Fig. 2h**). To further test whether mono-AMs might contribute to the *Lyz2*^Cre/+^;*Egr2*^fl/fl^ SiglecF^lo^ compartment, we examined the expression of *Cd93*, which we and others ^41^ have found to be progressively downregulated by monocyte-derived cells as they differentiate into AM. A stepwise decrease in *Cd93* expression throughout the integration trajectory was observed in *Egr2*^fl/fl^ controls (**Fig. 2p**). In contrast, while CD11b^+^ macrophages from *Lyz2*^Cre/+^;*Egr2*^flox/flox^ mice expressed high levels of *Cd93,* there was no progressive downregulation observed (**Fig. 2p**), but a sudden drop to the level observed in UV-RSV SiglecF^lo^ (of foetal origin). This pattern was not restricted to *Cd93.* In *Egr2*^fl/fl^ controls there was a reciprocal and stepwise increase in expression of *Fabp4*, a lipid chaperone important for surfactant recycling that is highly expressed in fAMs (**Fig. 2p**). This was not seen in *Lyz2*^Cre/+^;*Egr2*^fl/fl^ mice. This did not reflect EGR2-dependence of *Fabp4 per se*, as evidenced by equivalent expression of *Fabp4* in resident CD11b^−^ AM from *Lyz2*^Cre/+^;*Egr2*^fl/fl^ mice with controls (**Fig. 2p**). Thus, loss of CD11b^+^ mono-AM in *Lyz2*^Cre/+^;*Egr2*^flox/flox^ mice at 2mo post RSV most likely reflects cell attrition due to failure to integrate. Altogether, these results suggest that in the absence of EGR2, newly elicited CD11b^+^ macrophages are unable to increase their proliferative rates, fail to integrate and thus cannot contribute to the differentiated, self-renewing resident AM compartment.

### Developmentally discrete AM have distinct transcriptional and functional profiles

Our data above from *Egr2*^fl/fl^ controls suggested that RSV-elicited mono-AM that had integrated retain transcriptional differences relative to their fAM counterparts, long after viral clearance. However, our experiments in *Egr2*^fl/fl^ mice did not allow for definitive assessment of mono-AM and fAM. To test this directly, we used a variant of the tissue protected chimera system described above combined with RSV infection to faithfully identify foetally derived and bone marrow-derived AM (**Fig. 3a**). Specifically, recipient mice were administered the chemotherapeutic myeloablative agent busulfan to deplete host bone marrow without depleting AMs and other peripheral immune cell populations^29^. As this results in only partial repopulation with donor bone marrow, we exploited the CCR2-dependency of monocytes for egress from the bone marrow^42^ by using *Ccr2^−/–^* CD45.1^+^/CD45.2^+^ hosts. This results in >90% chimerism of donor CD45.2^+^ cells in the circulating monocyte pool (rather than the ∼40-50% seen in normal WT > WT busulfan chimeric mice)^29^ with essentially no chimerism in the AM compartment from busulfan chimeric mice with no other manipulation (**Fig. 3b**). After eight weeks of reconstitution, busulfan chimeric mice were inoculated with RSV and fAM (CD45.1^+^/CD45.2^+^) and elicited mono-AM (CD45.2^+^) purified by fluorescence activated cell sorting (FACS) at 1mo, 2mo and 4mo post-infection. Consistent with all other models above, mono-AM were present in RSV infected mice and their presence increased with time (**Fig. 3b, Extended data Fig. 3a**). We then cultured these cells overnight in medium alone, or with the TLR2 agonist Pam3CSK4, to investigate their responses to TLR ligation, and performed bulk RNA-seq.

**Fig. 3.**
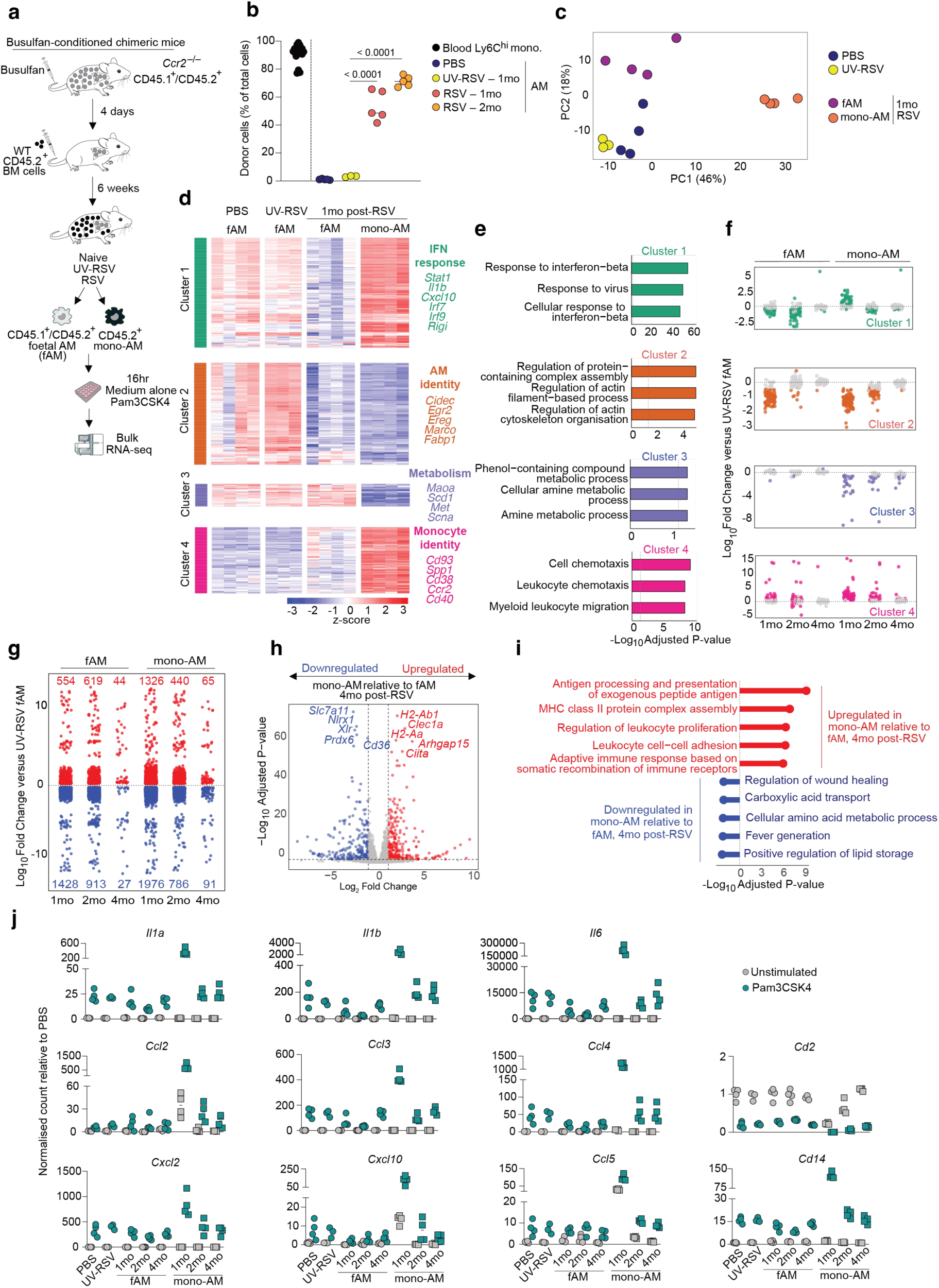
Developmentally discrete AM have distinct transcriptional profiles and trajectories. **a.** Experimental scheme for the generation of WT > *Ccr2^−/–^* busulfan chimeric mice, experimental groups and conditions for RNA-seq. **b.** Chimerism of CD11c^+^SiglecF^+^ AM in WT > *Ccr2^−/–^* busulfan chimeric mice inoculated with RSV 1 and 2 months prior or UV-RSV control. Symbols represent individual mice. Data from one experiment with *n* = 3-5 mice per group. Statistical significance calculated by one-way ANOVA. **c.** Principal component analysis (PCA) of bulk RNA-seq data from unstimulated AM subsets at one, two and four months after RSV or UV-RSV inoculation. **d.** Heatmap of hierarchically clustered top 500 most variable genes from bulk RNA-seq data of unstimulated treatment groups, annotated with selected cluster-defining genes. **e.** Pathway analysis (over-representation analysis) of genes within each cluster in (d). **f.** Log-fold change of each cluster gene relative to UV-RSV in fAM or mono-AM samples 1, 2, and 4mo post-RSV. Colouration denotes significance compared to UV-RSV fAMs. **g.** Counts of selected cluster-defining genes normalized to UV-RSV control, log1p transformed. Dotted line denoted average of UV-RSV. **h.** Log10 fold change of genes compared to PBS control. Significant genes denoted in red (upregulated) or blue (downregulated). The number of significant DEG shown. **i.** Over-representation analysis of genes upregulated and downregulated in mono-AM compared to fAM at 4mo post-RSV. **j.** RNA-seq data showing expression of cyto- and chemokine genes within indicated Pam3CSK4-stimulated or unstimulated AM subsets following PBS or UV-RSV, or one, two or four months after RSV inoculation. Points represent biological replicates.

Focussing on samples in the absence of TLR ligation, as expected, we found fAM from naïve and UV-RSV chimeric mice were transcriptionally almost identical, confirming the minimal effects of UV-RSV (**Fig. 3c, d**). Clustering analysis confirmed clear transcriptional differences between fAM and mono-AM post-RSV and identified 4 modules of genes that had distinct expression patterns across subsets and time (**Fig. 3d**), for simplicity referred to as “IFN response”, “AM identity”, “metabolism” and “monocyte identity” clusters respectively. For instance, genes in the IFN response cluster, which were associated with anti-viral responses and interferon-associated pathways (e.g., *Stat1*, *Irf7* and *Cxcl10*), were more highly expressed by mono-AM at 1mo post RSV than any other group (**Fig. 3d-f**). Strikingly, fAM in the post-RSV (1mo) lung expressed IFN response cluster genes at a lower level than their counterparts in control groups (naïve and UV-RSV) (**Fig. 3d-f**). By 2mo post RSV, these genes were further downregulated in fAM, suggesting sustained downregulation of this gene module in fAM in the post-RSV lung (Fig. 3f). Notably, there was marked downregulation of these genes in mono-AM at 2mo post-RSV compared to fAM in control mice, suggesting that the lung environment drives this effect on macrophages independently of their ontogeny, but with a temporal delay in the mono-AM compartment (**Fig. 3f**). Genes in the AM identity cluster were associated with cytoskeletal organization machinery (e.g., *Rhoc, Arpc4*) and contained many genes considered to be part of the homeostatic AM signature (e.g., *Krt19, Cidec, Marco, Egr2, Fabp1*). Expression of genes in the AM identity cluster was profoundly downregulated in fAMs at 1mo post RSV but returned to baseline levels by 2mo post RSV (**Fig. 3d-f**). By contrast, AM identity cluster genes were expressed at low levels in mono-AM at 1mo post-RSV but required 4 months to return to baseline, consistent with mono-AM slowly but progressively adopting the identity of their fAM counterparts. Genes forming the metabolism cluster were mainly associated with metabolism (e.g., *Scd1*, *Slc7a11*, *Lepr, Plcb1*) and remained unchanged in fAMs post-RSV when compared with control groups, whereas mono-AM acquired expression of these genes over time, although these never reached the same level as in fAM (**Fig. 3f**). Finally, many genes in the monocyte identity cluster were associated with chemotaxis (e.g., *Ccr1, Ccr2*) and proinflammatory effector function (e.g., *Ccl2, Ccl7, Cd40, Spp1*) and were highly expressed by mono-AM at 1mo post-RSV, with minimal changes in expression within fAMs. Consistent with progressive maturation of mono-AM, expression of monocyte identity cluster was lower at 2mo post-RSV compared with 1mo (**Fig. 3d-f**). Thus, gene expression changes post-RSV are a function of AM subset, gene module, and time.

To identify long-term, persistent gene expression differences post-RSV, we performed untargeted DEG analysis of mono-AM and fAM at 4mo post RSV. While there was convergence in transcriptional signatures, differences were still apparent (**Fig. 3g-i**). Differential gene expression and pathway analysis demonstrated that many genes involved in antigen processing and presentation (*H2-Ab1, H2Aa, Ciita*) were more highly expressed in mono-AM compared with fAM (**Fig. 3h-i**). Amongst fAM, expression of *Slc7a11* and *Nlrx1* were some of the most DEG, which have been associated with control of metabolic processes. This was confirmed by over-representation pathway analysis (**Fig. 3i**). These observations highlight that mono-AM retain transcriptional differences for prolonged periods, particularly as relates to MHCII and metabolic gene expression profiles.

To determine if functional differences were present between AM subsets post RSV, we next assessed the responsiveness of the developmentally distinct AMs to stimulation. As expected, treatment with the TLR2 agonist Pam3CSK4 treatment resulted in major transcriptional changes (**Fig. 3j**). Again, AM from control groups (naïve and UV-RSV) exhibited comparable transcriptional responses. In contrast, mono-AM from 1mo post RSV displayed the largest degree of transcriptional change, with profound transcriptional upregulation of multiple cytokine and chemokine genes, including *Il1a, Il1b, Il6, Cxcl10, Ccl2, Ccl3, Ccl4, Ccl5,* exceeding the response observed in fAM in the lung (**Fig. 3j**). This suggests that fAM and mono-AM, despite residing in the same environment, respond differently to identical stimuli, with sustained TLR hyperresponsiveness of mono-AM post RSV. Intriguingly, for a subset of genes including *Il1b, Il6, and Ccl3,* fAM showed reduced responsiveness post-infection compared with pre-infection, indicating that resident macrophages may exhibit partial hypo-responsiveness post-infection (**Fig. 3j**). In some instances, the fAM responsiveness is even more diminished at 2mo compared with 1mo post-RSV (*Ccl5*, *Cxcl10*) (**Fig. 3j**). Taken together, these analyses show that RSV infection results in both short- and long-term transcriptional changes to the AM compartment. Distinct AM subsets show discrete molecular reprogramming, characterized by hyperresponsiveness and delayed metabolic adaptation within mono-AMs, while fAMs show minimal or reduced stimulus responsiveness and an attenuated interferon response signature.

### Outcompetition and divergent transcriptional profiles are conserved features of mono-AMs independent of infection

The phenotypic identity and behaviour of mono-AM in the context of RSV was reminiscent of that observed in other infection and challenge contexts in the lung, particularly IAV^29, 35^. This prompted us to examine whether these features are virally-induced or whether there may be generalizable phenotypic and functional features of monocyte-derived macrophages in the lung, independent of the eliciting challenge. Thus, we next employed a model to elicit monocyte-derived macrophages in an infection-independent context using clodr-lipo administration and compared our findings to those observed in RSV and IAV infection as a second, distinct viral infection context. When administered intratracheally to the bronchoalveolar space, efficient phagocytosis of clodr-lipo by AM results in high intracellular concentrations of clodronate, which is metabolised intracellularly to an ATP analogue that competitively inhibits the ADP/ATP translocase and is therefore cytotoxic, resulting in AM cell death^43^. Two administrations of clodr-lipo resulted in depletion of AM numbers to around 10% of the steady state numbers 3 days after administration (**Extended Data Fig. 4a, b**), with the AM pool steadily repopulating over the weeks following administration (**Extended Data Fig. 4c**). Multiplexed cyto-/chemokine analysis of BALF supernatants show that clodr-lipo cause minimal inflammation compared with RSV or IAV infections **(Fig. 4a)**. Furthermore, barrier integrity was not compromised by clodronate treatment, as red blood cell (RBC) and albumin levels did not significantly increase after clodr-lipo administration, unlike in RSV and IAV (**Fig. 4b**). Mice treated with clodr-lipo also did not exhibit weight loss, similar to RSV and in contrast to IAV infection (**Extended Data Fig 4d**). Histological analysis revealed little inflammation following clodr-lipo, with changes limited to local cellular infiltration around the vasculature at early timepoints (**Extended Data Fig. 4e, f**). Nevertheless, monocytes were recruited that gave rise to a population of CD11b^+^ macrophages, which peaked at 14d, but had contracted by 18d and were essentially absent by 1mo after clodr-lipo administration **(Fig. 4c, d)**. We also noted the emergence of a more stable population of SiglecF^lo^ CD11c^+^ cells **(Fig. 4c, d)**. Given our observations and those of others that mono-AM express less SiglecF than fAMs^35^, this indicated that mono-AM were being generated by clodr-lipo administration. Consistent with this, SiglecF^lo^ CD11c^+^ macrophages in this model expressed higher levels of CD93 compared with their SiglecF^hi^ counterparts (**Fig. 4e**). By using the *Ms4a3*^Cre/+^*;Rosa26*^CAG-LSL-tdTomato/+^ system, we showed these SiglecF^lo^ AMs were labelled to a high level, confirming their monocyte origin **(Fig. 4f**). As in RSV, the tdTomato^+^ cell fraction in the total AM pool increased over time, consistent with progressive displacement of fAM by mono-AM (**Fig. 4g**).

**Fig. 4.**
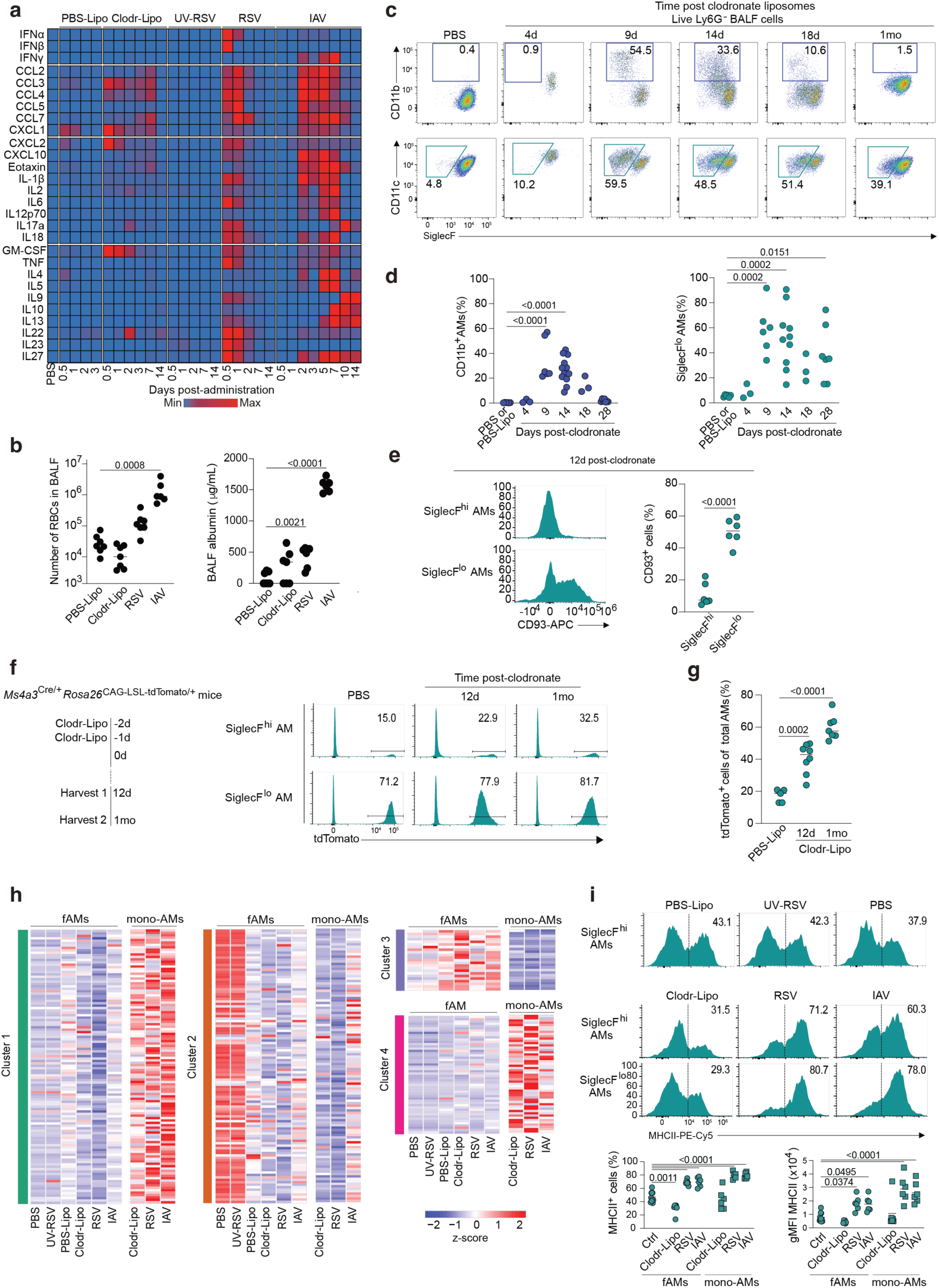
Outcompetition and divergent transcriptional profiles are conserved features of mono-AMs independent of infection. **a.** Heatmap showing cyto-/chemokine levels in bronchoalveolar lavage fluid of mice at indicated timepoints after clodr-lipo administration, RSV or IAV (1×10^4^ TCID_50_) infections. Columns represent averaged value across mice from 7 independent experiments of *n* = 3-5 per timepoint per condition. Colours represent normalized levels within an analyte across conditions and timepoints. **b.** Red blood cell (RBC) number (left) and albumin content (right) of BALF two days after clodr-lipo administration or –RSV or 7 days post-IAV reflecting the peak of damage. Symbols represent an individual mice. Data pooled from two independent experiments with *n* = 6-7 mice per group. Significance calculated by one-way ANOVA. **c.** Representative flow cytometry plots of and CD11b versus SiglecF (top) and CD11c versus SiglecF (bottom) by Live, Ly6G^-^ BAL cells obtained from naive C57BL/6 mice or mice administered clodr-lipo and then harvested at the indicated time points. **d.** Quantification of CD11b^+^ (left) and SiglecF^lo^ (right) AM at indicated timepoints after clodr-lipo administration. Data pooled from 4 independent experiments. Symbols represent individual mice from *n* = 3-10 mice per timepoint. Statistical significance calculated by one-way ANOVA. **e.** Representative histograms (left) and quantification (right) of CD93 expression by SiglecF-defined macrophage subsets at 12d after clodr-lipo administration. Each point (right) represents one biological replicate. Statistical significance calculated by two-tailed *t*-test. **f.** Experimental scheme for clodr-lipo administration in *Ms4a3*^Cre/+^;*Rosa26*^CAG-LSL-tdTomato/+^ mice (left) and representative histograms showing tdTomato expression by SiglecF^hi^ and SiglecF^lo^ AM subsets from *Ms4a3*^Cre/+^;*Rosa26*^LSL/CAG-tdTomato/+^ mice harvested at the indicated time points post clodr-lipo administration (right). **g.** Frequency of tdTomato positive cells within SiglecF^hi^ and SiglecF^lo^ AM subsets (pre-gated as Live MerTK^+^ CD64^+^SiglecF^+^CD11c^+^) obtained from *Ms4a3*^Cre/+^;*Rosa26*^CAG-LSL-tdTomato/+^ mice harvested at the indicated time points after clodr-lipo administration liposome administration. Symbols represent individual mice from *n*=5-8 mice per timepoint. Statistical significance calculated by one-way ANOVA. **h.** Heatmap showing expression levels of gene clusters defined in Fig. 3d. Clodronate samples collected at 0.5mo, RSV and IAV samples collected at 1mo. Normalized to PBS control of each experiment, z-score plotted. Average of samples within each condition displayed. **i.** Representative histograms (top) and quantification of frequency of positive cells and gMFI (bottom) for MHCII within SiglecF^lo^ and SiglecF^hi^ AM subpopulations (pre-gated as Live MerTK^+^ CD64^+^SiglecF^+^CD11c^+^) at 12d post-PBS lipo (control), 12d post clodronate, 1mo RSV, 1mo UV-RSV (control), 1mo IAV and 1mo PBS (control). Symbols represent individual mice. Data from one experiment with *n* = 4-6. Statistical significance calculated by one-way ANOVA.

Unlike RSV infection, monocytes elicited by clodr-lipo treatment arrive into, and differentiate within, an environment free of infection. We leveraged this system to dissect aspects of monocyte differentiation specific to viral infection from those that represent a hard-wired process independent of infection. Using our busulfan chimeric mouse model (**Fig. 3a**), we isolated fAMs and mono-AMs from mice after clodr-lipo administration or IAV infection, to compare transcriptomes of these AM subpopulations across eliciting contexts. First, we used the gene clusters defined in our analysis of RSV, which represented the most differentially expressed genes between fAM and mono-AM, to determine the relative overlap across contexts. To our surprise, clodronate treatment largely recapitulated the transcriptional differences across the different gene modules. Indeed, there was a high degree of conservation in the gene expression in mono-AM across clodronate, RSV and IAV, indicating that many of the features of mono-AM seen in our RSV dataset mainly represent hard-wired features of monocyte differentiation and not response to viral infection, even including typical ‘anti-viral’ features in the IFN response cluster, such as *Cxcl9* and *Cxcl10* expression (**Fig. 4h**).

Not all transcriptional features of mono-AMs were conserved across eliciting contexts; our transcriptional analysis highlighted a sustained upregulation of pathways involved in antigen processing and presentation in mono-AM compared with fAM, even at 4mo post-RSV **(Fig. 3h, i**). However, while we found that MHCII expression was elevated on the surface of both fAM and mono-AM after both viral infections, this was not the case following clodr-lipo administration (**Fig. 4i**), suggesting that a post-infectious lung is required for MHCII expression in mono-AMs. Further comparison within the subsets showed a slightly higher expression of MHCII within mono-AMs compared with fAMs after viral infection (**Fig 4i**), indicating that cell origin interacts with the environment to determine the level of MHCII. Together, these results suggest that the transcriptional profile of mono-AMs is largely conserved across contexts, but that some reatures some features of AM after infection, such as elevated MHCII, may be contributed by the lung environment.

### Mono-AM immunoreactivity is independent of recruitment context and contributes to anti-bacterial protection

Our comparative transcriptional analysis suggested marked conservation of the monocyte integration and differentiation process across eliciting contexts. Thus, we compared side-by-side the functional features of mono-AMs across RSV, IAV and clodr-lipo to assess whether and how differences in infection and inflammation status might impact on mono-AM functionality. First, we sought to determine whether protracted metabolic adaptation in the bronchoalveolar space was a hardwired feature of monocyte-derived cells. As lipid metabolism is a key part of homeostatic AM function, we measured uptake of the fluorescently labelled long-chain fatty acid (FA) BODIPY-C16 by flow cytometry and found that mono-AMs across the board showed equivalent significantly reduced capacity to take up this molecule (**Fig. 5a**). This did not represent a global inability to uptake nutrients, as uptake of the fluorescent glucose analogue 2-NBDG was equivalent between fAM and mono-AM (**Extended Data Fig. 5a**). Given the role of peroxisomes in branched chain and very long chain FA metabolism, we measured peroxisome mass and found that mono-AM failed to achieve the high level detected in fAMs (**Fig. 5b**). A similar finding was seen with mitochondrial mass (**Extended Data Fig. 5b**). Given the reduction in mitochondrial mass, we investigated mitochondrial dependency for ATP production by profiling fAM and mono-AM using the SCENITH (Single Cell ENergetIc metabolism by profiling Translation inHibition) assay^44^. SCENITH exploits the fact that around half of the energy generated in mammalian cells is immediately consumed by protein synthesis processes. Thus, by measuring the rate of protein synthesis through detecting incorporation of the drug puromycin, in the presence metabolic inhibitors alone or in combination (oligomycin and 2-deoxy-D-glucose), metabolic profiling can be combined with advanced immunophenotyping. As expected, fAM across contexts showed near complete mitochondrial dependence that was not altered by infection. By contrast, elicited mono-AMs across contexts exhibited significantly reduced mitochondrial dependence (**Fig. 5c**) compared with fAMs from both infected and naïve controls. Both subsets showed high glucose dependence across contexts, indicating that, whether directly through glycolysis or through OXPHOS, both subsets highly depend on glucose to meet their energetic needs (**Fig 5c**). Using transcriptional profiling, lipid uptake, and SCENITH analysis, these results highlight that mono-AMs do not display the metabolic profile of their resident counterparts, despite residing in the same bronchoalveolar environment for nearly a month.

**Fig. 5.**
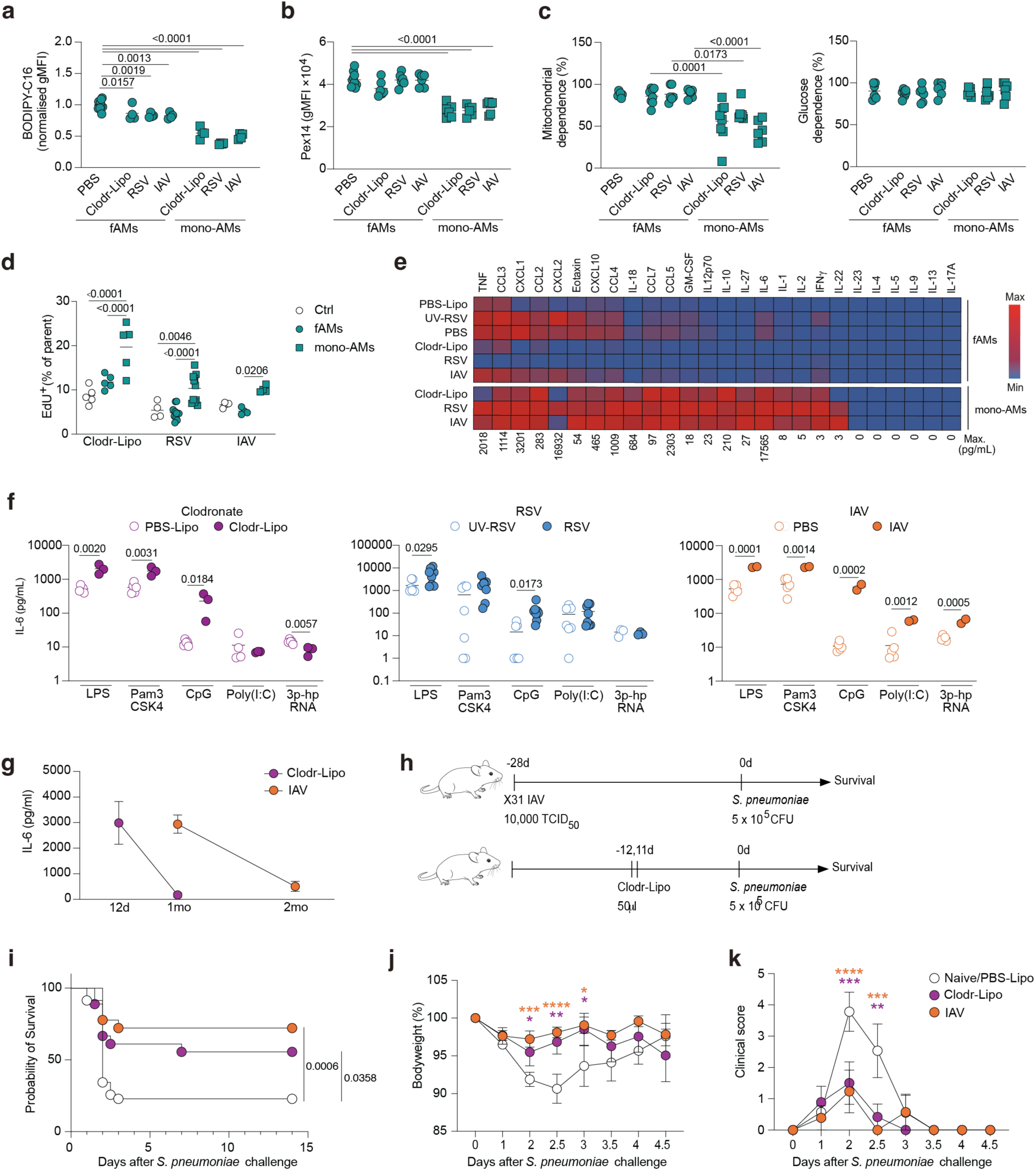
Metabolic, proliferative and immunoreactivity profiles of monocyte-derived AMs are conserved across contexts and can drive *S. pneumoniae* protection in the absence of prior infection. Mice were infected with 5×10^5^ PFU RSV, 10,000 TCID_50_ X31 or treated twice with 50μl clodr-lipo intratracheally. **a.** Uptake of long-chain fluorescent fatty acid BODIPY-C16 by AMs *in-vitro* from 12d after clodr-lipo administration, 1mo post-RSV and 1mo post-IAV mice. AM were separated on origin using SiglecF expression. Symbols represent biological replicates. Data pooled from three representative experiments of six, with *n* = 4 mice per group. Significance assessed by one-way ANOVA. **b.** Abundance of PEX14 protein as a proxy of peroxisome mass in AM at 12d after clodr-lipo administration, 1mo post-RSV and 1mo post-IAV. AM were separated on origin using SiglecF expression. Symbols represent biological replicates. Data from one representative experiment of four with *n* = 4-6 mice per group. Significance assessed by one-way ANOVA. **c.** SCENITH analysis of AMs at 12d after clodr-lipo administration, 1mo post-RSV and 1mo post-IAV. AM were separated on origin using SiglecF expression. Each point represents one mouse with *n* = 6-8 mice per timepoint. Data pooled from 6 independent experiments. Significance assessed by one-way ANOVA with pairwise comparisons to the naive condition. **d.** EdU incorporation into AM at 12d after clodr-lipo administration (ctrl = 12d PBS-Lipo), 1mo post-RSV (ctrl = 1mo UV-RSV) and 1mo post-IAV (ctrl = 1mo PBS) at 24 hours after EdU injection. AMs were separated on origin using SiglecF expression. Significance assessed by two-way ANOVA with pairwise comparison of AM subsets within experimental groups (Cl.-Lipo, RSV, IAV). Data pooled from four independent experiments. *n* = 4-11 per group. **e.** Cyto- and chemokine array of supernatants from fAM and mono-AM from busulfan chimeras, FACS sorted using congenic markers within SiglecF^+^ CD11c^+^ AM and *ex-vivo* stimulated overnight with 100ng/mL Pam3CSK4. Heat map represents normalized values across conditions within an analyte. Data pooled from two independent experiments with *n* = 3-5 mice per condition. **f.** IL-6 production assessed by ELISA from bulk AM at 12d after clodr-lipo administration or –PBS liposomes, 1mo post-RSV or –UV-RSV, and 1mo post-IAV and naive mice stimulated overnight in the presence of 100ng/mL LPS, 100ng/mL Pam3CSK4, 1uM CpG, 50ug/mL Poly(I:C), 400ng/mL 3p-hpRNA. Points represent individual mice. Significance assessed by multiple T-tests comparing within each TLR agonist between experimental and control conditions. Data from 3 representative of 6 independent experiments with *n* = 2-7 mice per group. Significance calculated by multiple unpaired *t* tests. **g.** IL-6 production over time by bulk AM from after clodr-lipo administration and post-IAV mice at indicated timepoints. **h.** Schematic of survival experiment. Mice were infected with influenza 1 month, or treated with clodr-lipo 12 days, prior to infection with 5×10^5^ CFU *S. pneumoniae* (TIGR4). **i.** Kaplan-Meier plot representing cumulative survival of mice after *S. pneumoniae* challenge. Data from n=18-35 mice per group. Data plotted as mean±SEM. Significance assessed by log-rank (Mantel-Cox) test. **j.** Weight loss of mice at indicated timepoints following *S. pneumoniae* infection. Mice exceeding humane endpoints were removed from weight loss and clinical scores in later timepoints. Data from *n* = 18-35 mice per group. Data plotted as mean ±SEM. Significance assessed by two-way ANOVA comparing groups within timepoints. Asterisks represent \**P*<0.05, \*\**P*<0.01, \*\*\**P*<0.001, \*\*\*\**P*<0.0001 **k.** Clinical scores of mice at indicated timepoints following *S. pneumoniae* infection. Mice exceeding humane endpoints were removed from weight loss and clinical scores in later timepoints. Data from *n* = 18-35 mice per group. Data plotted as mean ±SEM. Significance assessed by two-way ANOVA comparing groups within timepoints. Asterisks represent \**P*<0.05, \*\**P*<0.01, \*\*\**P*<0.001, \*\*\*\**P*<0.0001

Next, we asked whether enhanced proliferative activity was a conserved feature across contexts. EdU incorporation confirmed that, across the contexts, mono-AM displayed enhanced proliferative rates compared with fAM (**Fig. 5d**). Similar results were obtained with Ki67 staining (**Extended Data Fig. 5c**). Slightly higher overall levels of proliferation were observed after clodr-lipo administration, likely due to recent/ongoing repopulation at this timepoint (**Fig. 4d**). Thus, enhanced proliferation rate is a conserved feature of mono-AMs, likely underpinning their ability to outcompete their resident counterparts across environmental contexts.

Another defining feature of mono-AMs compared to fAMs in post-viral environments is their altered responsiveness to TLR stimulation, a feature often assumed to be driven by the altered tissue environment^45^. To determine if this is the case, we again performed a comparative analysis of responsiveness across eliciting contexts. Strikingly, mono-AM elicited by clodr-lipo displayed the same hyperresponsiveness to TLR (Pam3CSK4) stimulation seen in mono-AM elicited by RSV or IAV. Mono-AM elicited by clodr-lipo not only were hyperresponsive, but had an almost identical secretome to that of mono-AM elicited by RSV or IAV (1mo post infection) as assessed by cyto-/chemokine array (**Fig. 5e**). By contrast, across contexts, fAM were relatively refractory to TLR stimulation (**Fig. 5e**), particularly following RSV infection, reflecting changes seen at transcriptional level (**Fig. 3j**). These results suggest that fAM and mono-AM mount divergent responses to identical challenge, with mono-AM displaying high immunoreactivity and fAM consistently remaining hyporesponsive.

To determine the aggregate response of the AM population to a greater breadth of potential stimuli, we isolated total AMs from BALF following clodronate, RSV and IAV and assessed the responsiveness to a variety of pattern recognition receptor (PRR) agonists. IL-6 was used as a sensitive and reliable readout based on our multiplex analysis (**Fig. 5e**) and our previous work^29^. Across contexts and stimuli, higher levels of IL-6 were detected following challenge, demonstrating that the AM population on aggregate is hyperresponsive, and that compositional changes have broad functional impacts (**Fig. 5f**). Importantly, the frequency of mono-AMs amongst the total BALF AM population was not significantly different across recruitment contexts (**Extended Data Fig. 5d**). In the context of IAV, hyperresponsiveness was observed following the delivery of each stimulus. In contrast, for RSV and clodr-lipo, similar responses were seen to that of IAV following LPS and CpG stimulation, but not Poly(I:C) or 3p-hpRNA (**Fig. 5f**).

Furthermore, despite the similarities in responsiveness by mono-AM across contexts, there were clear differences in the rate at which monocyte-derived cells integrated and the associated hyperresponsiveness waned. Again, using IL-6 production as a readout of responsiveness, we assessed mono-AM longitudinally following clodronate or IAV, the systems representing the least and the most inflammatory of our models. While IL-6 production by clodronate-elicited mono-AM had returned to baseline levels seen in fAM counterparts by 1mo, waning within the post-IAV environment took much longer, with IL-6 production only nearing baseline levels by 2mo post-infection (**Fig. 5g**). Thus, heightened responsiveness is a feature of monocyte-derived cells but the environment in which these cells integrate impacts the speed of functional adaptation.

Finally, to dissect the consequences of ontogeny and environmentally controlled features in subsequent immune challenge, we performed a secondary bacterial infection with *Streptococcus pneumoniae*. Mice were infected with *S. pneumoniae* at 12d after clodr-lipo administration or 1mo post-IAV infection, the point at which maximal hyperresponsiveness was detected (**Fig. 5h**). Strikingly, both IAV- and clodronate-experienced mice displayed improved survival outcomes to *S. pneumoniae* infection (**Fig. 5i**), with concomitant reduction in both weight loss (**Fig. 5j**) and clinical scores (**Fig. 5k**). While it was previously shown that IAV-elicited mono-AM control subsequent *S. pneumoniae*^29^, these results highlight that the superior role of mono-AMs in clearing bacterial infection represents hardwired features of monocyte-derived AMs rather than imprinting by the post-viral lung environment. Together, our results demonstrate that monocyte ontogeny predominantly drives the functional profile of mono-AMs, but that the lung environment can further fine-tune their features qualitatively and over time.

## Discussion

Residing in the alveolar space, AMs are frequently a first target of invading pathogens and insults. Here, we show that, similar to other viral infection contexts, RSV infection results in partial depletion of the fAM compartment and the appearance of mono-AMs. Through comparative analysis across infectious and sterile recruitment contexts, we reveal that monocyte differentiation in the bronchoalveolar space follows a conserved blueprint irrespective of eliciting agent which is characterized by enhanced proliferation, reduced mitochondrial metabolism, enhanced production of a fixed set of cytokines and chemokines, and enhanced sensitivity to TLR stimulation. Collectively, our findings challenge the notion of inflammation- or infection-driven macrophage differentiation and rather suggests that the developmental trajectory and functionality of mono-AM is hard-wired and determined by their prior monocyte identity, a feature that we have previously termed monocyte legacy^46^. This hard-wired, conserved trajectory can be further fine-tuned by the lung environment.

Our study builds on previous work that focussed on the acute phase of RSV infection and was performed at a time when distinction between macrophages, monocytes and differentiation intermediates as well as genetic lineage tracing tools were still in their infancy^39,47^. Through longitudinal studies in mice that include lineage identification, we show that the lung steady-state, where AM niches are filled by fAMs with little contribution of recruited bone-marrow derived cells, is perturbed by infection-induced disappearance of fAMs through their decreased proliferation and increased apoptosis, which frees up niches for a biphasic repopulation: Initially through proliferation of surviving fAMs, followed by their displacement by monocyte-derived cells. Once the available lung niches are filled, there is no further recruitment into the lung, and the lung becomes a largely closed environment again.

We found that mono-AM differentiate during engraftment from a CD11b^+^ to a CD11b^—^state. This transition was underpinned principally by the acquisition of supra-homeostatic proliferative capacity, dependent on the transcription factor EGR2. Given their prolonged tissue residency and relative independence from circulation for maintenance, self-renewal is a key property of tissue-resident macrophages. In *Lyz2*^Cre/+^;*Egr2*^fl/fl^ mice, CD11b^+^ cells were seemingly unable to transition into the self-renewing resident AM compartment. Counterintuitively, fAM in *Lyz2*^Cre/+^;*Egr2*^fl/fl^ mice are able to survive in the lung long-term, even repopulating the lungs after lung insult, which suggests relatively intact self-renewal capacity. The discrepancy in phenotype may thus relate to precursor characteristics. Previous work comparing yolk-sac primitive macrophages, foetal liver monocytes and bone marrow monocytes reveals that proliferation is one of the most significantly enriched pathways in foetal progenitors compared with adult bone marrow monocytes^48^. Therefore, it is possible that such previously-endowed proliferative capacity may be sufficient to overcome EGR2 deficiency in the embryonic, but not the adult HSC-derived, monocyte compartment.

Our results reveal an increased responsiveness of AMs on the population level following engraftment of highly immunoreactive mono-AMs. This phenomenon is distinct to trained immunity, where the same cellular population or their direct offspring acquire memory-like properties upon exposure to cytokines or pathogen associated molecular patterns. This exposure triggers metabolic or epigenetic rewiring that renders cells hyperresponsive to secondary stimulation^49^. Recent studies have identified roles for peripheral trained immunity in reprogramming functionality of fAMs after insult^50, 51^. While both trained immunity and recruitment of immunoreactive mono-AM will have the effect of causing hyperresponsiveness on a population level, these represent two fundamentally distinct phenomena at a cellular and mechanistic level. We observe that pre-existing fAMs show only limited transcriptional and functional changes upon infection; instead, mono-AMs emerging after infection display distinct functionality, likely as a result of their blood monocyte origin. Distinguishing these phenomena will be key for therapeutic intervention as they are likely to be driven by distinct biological mechanisms.

The primary function of AM during homeostasis is clearance of surfactant, a process which relies on intact lipid uptake and catabolism. The two key organelles involved in lipid metabolism are peroxisomes and mitochondria. Peroxisomes play a key role in branched-chain and very long-chain fatty acid metabolism^52^ and are essential for proper homeostatic and immune function of AM^53, 54^. Mitochondria are also critical for fatty acid β-oxidation and for the utilization of the resulting reduced cofactors and acetyl-CoA for the production of ATP via the TCA cycle. In addition to superior proliferative capacity and immunoreactivity, mono-AM exhibit a divergent metabolic profile, characterised by lower peroxisome and mitochondrial mass compared with fAM. Previous studies have shown that, in the absence of functional peroxisomes, AM showed impaired mitochondrial health, lipotoxicity and cell death, which led to impaired epithelial repair and enhanced mortality in the context of viral infection^53, 54^. The reduced abundance of peroxisomes in mono-AM may thus impact on their function in the context of heterologous infection; indeed, the presence of mono-AM is pathogenic during IAV infection^35^. Furthermore, mono-AM had reduced mitochondrial mass and reduced uptake of long-chain fatty acids. Given these observations, we assessed whether there were differences in mitochondrial dependence for ATP production between the subsets. It has been previously shown that AM are highly reliant on OXPHOS to process lipids; indeed, loss of *Tfam*, which controls both replication and transcription of mitochondrial DNA, in the AM compartment results in pulmonary alveolar proteinosis^55^. Using SCENITH analysis, we found that mono-AMs show a significantly reduced mitochondrial dependency across contexts compared with their fAM counterparts. Overall, these origin-dependent metabolic characteristics appear to be largely independent of the lung environment as they persist for extended periods of time with very slow adaptation. This may affect long-term the uptake and turnover of lipids in the lung, including surfactants, after infection, with potential implications for the return to full lung functionality.

The link between glycolysis and proliferation is well-known; glycolysis acts as a central node for several anabolic/biosynthetic pathways, such as the pentose phosphate and serine synthesis pathways, which generate nucleotides, amino acids and phospholipids required for proliferation^56^. We speculate that the dependence on glycolysis, likely inherited from their blood monocyte precursors, may provide mono-AM with a metabolic predisposition that allows for more rapid proliferation. Over time, this superior proliferative capacity leads to complete replacement of the fAM compartment. How such replacement dynamics would play out in the context of a second infection has not been explored but would likely be dependent on the speed of metabolic and proliferative adaptation of the first wave of mono-AM.

Not all RSV-induced changes in the AM compartment were related to origin. A cluster of genes including AM identity genes was downregulated in both fAM and mono-AM after RSV infection, which was enriched in pathways relating to the chemotaxis and the actin cytoskeleton and also contained the transcript for the gene encoding the transcription factor EGR2. Short term, reversible loss of AM identity gene expression within fAMs has also been observed after IL-33 administration to mice^57^, suggesting this may be a response to acute inflammation. Furthermore, we observed prolonged elevated expression of MHCII in post-RSV fAM, as well as mono-AM by flow cytometry, as has previously been reported in lung mononuclear phagocytes^58^. By contrast, this was not the case in fAM and mono-AM after clodr-lipo administration. Respiratory viral infection is succeeded by the establishment of tissue-resident memory T cells, which does not occur after sterile depletion with clodr-lipo. Therefore, it is unsurprising, and likely, that signals such as IFN-γ, a known inducer of MHCII, from such tissue-resident lymphocytes after infection may result in the upregulation of such molecules on AM after infection. Indeed, depletion of T cells in post-IAV lungs attenuates infection-induced MHCII upregulation^51^.

We also observed differences in PRR agonist sensitivity between clodr-lipo, RSV and IAV-experienced AMs. While post-IAV AM showed enhanced IL-6 production in response to LPS, Pam3CSK4, CpG, Poly(I:C) and 3p-hpRNA stimulus, PRR-sensitivity of after clodr-lipo administration and –RSV AMs was restricted to LPS, Pam3CSK4 and CpG. This suggests that signals during or following IAV infection can act on AM to modulate their PRR sensitivity. Interestingly, the expression of TLR9 is normally repressed in AM by the transcription factor KLF4, to limit inflammation caused by DNA sensing during steady state efferocytosis^19^. This suggests an enhanced sensitivity of mono-AM to dead cells, which may have implications for heterologous challenges and chronic inflammatory diseases. Together, these results suggests that, beyond monocyte legacy, viral infection can further modulate the profile of AM in an origin-independent manner due to infection-induced, cell-extrinsic changes to the lung environment.

Our transcriptional data revealed divergent gene expression trajectories of post-infection fAM and mono-AM with regard to IFN response cluster genes, comprised of interferon response and interferon signalling genes. While after RSV, expression of this subset of genes was elevated in mono-AM, it was reduced in fAM. A similar, albeit less pronounced, observation was made in clodronate-experienced fAM, while IAV-experienced fAM did not downregulate genes within this subset. Similar observations were made when comparing cyto-and chemokine secretion by fAMs after Pam3CSK4 stimulus across eliciting contexts. It is possible that the magnitude of depletion induced by the different insults may contribute to the frequency of surviving fAM which were exposed to insult; RSV induces significantly less depletion compared with IAV^35^. Thus, surviving fAM post-IAVmay be those that are most distal to the infection foci and therefore are less exposed to infection-related imprints. Furthermore, given that the proportions of mono-AMs versus fAMs is similar across the three insults, it is possible that post-IAV fAM require more proliferative cycles to repopulate to the same numbers, during which imprints may be lost.

Transcriptional and functional differences between mono-AMs and fAM were transient; by 4 months post-RSV, only 156 genes were differentially expressed between mono-AM and naive fAM. Furthermore, differences in cytokine production were lost at two months post-RSV, a process which has previously been described in the context of IAV infection as “immunosedation”^46^. This suggests that RSV-elicited mono-AM acquire an fAM signature after prolonged residence in the lung. The kinetic of immunosedation post-RSV emulates that previously described for IAV infection^29^. By contrast, in the context of clodr-lipo, loss of enhanced IL-6 production occurs faster, being fully attenuated at 1 month after clodr-lipo administration. This suggests that the process of immunosedation is tunable and sensitive to the lung environment, and the mechanisms underlying this may provide scope for therapeutic intervention in cases where persistent immunoreactivity underlies disease pathology.

At timepoints of elevated immunoreactivity, mono-AM across all recruitment contexts showed near-identical cytokine and chemokine secretion profiles upon stimulus with Pam3CSK4. This suggests that mono-AM recruited in these divergent contexts would have convergent functions in an infection challenge. Indeed, we observed that clodronate-elicited mono-AM were protective during *S. pneumoniae* infection, indicating that the presence of mono-AMs alone is sufficient to alter the outcome of subsequent infection, in the absence of prior infection. While protective in this context, others have observed that the presence of mono-AM can negatively affect the outcome of influenza infection and fibrosis^31, 35^. Our results suggest that origin alone could be sufficient to explain heterologous protection or sensitivity to subsequent lung insults, paving the way for a more generalizable interpretation and application of results uncovered in such studies.

Together, our results uncover that mono-AM share largely conserved functional properties, irrespective of the nature of the insult eliciting them. Thus, understanding conserved mechanisms regulating mono-AM function could have therapeutic implications for diverse lung pathologies wherein mono-AM are elicited.

## Materials and Methods

### Mouse lines

All mice were used on a C57BL/6 background and a mix of male and female mice were used for each experiment. All mice were bred and maintained at the BVS LFR facility (University of Edinburgh), BVS CRM facility (University of Edinburgh) or Francis Crick institute under specific pathogen free conditions. All procedures were approved by the UK Home Office and performed under PPLs P4871232F, PP4544912, PP7469641, PP1902420, PP9277301, P9C468066 and P2394CA47. Transgenic mice used in this study are listed in Table 1.

**Table 1.** Mouse strains used in this study, their source and relevant identifiers.

| Strain | Source | Publication |
| --- | --- | --- |
| C57BL/6J | Charles River |  |
| C57BL/6J CD45.1<sup>+/+</sup> | University of Edinburgh<br>The Francis Crick Institute |  |
| C57BL/6J CD45.2<sup>+/+</sup> | University of Edinburgh<br>The Francis Crick Institute |  |
| C57BL/6J CD45.1<sup>+</sup>/CD45.2<sup>+</sup> | University of Edinburgh<br>The Francis Crick Institute |  |
| *Lyz2*<sup>Cre/+</sup>; *Egr2*<sup>flox/flox</sup> | University of Edinburgh<br>The Francis Crick Institute<br>*Egr2*<sup>flox/flox</sup> line originally obtained from P.Wang, Queen Mary University London. | (McCowan et al., 2021) |
| *Ms4a3*<sup>Cre</sup>; *Rosa26*<sup>CAG-LSL-tdTomato</sup> | Generated by F. Ginhoux and sourced from R. Gentek, University of Edinburgh | (Liu et al., 2019) |
| *Ccr2*<sup>Cre-ERT2</sup>; *Rosa26*<sup>CAG-LSL-tdTomato</sup> | Generated by B. Becher and sourced from D. Zaiss, University of Edinburgh | (Croxford et al., 2015) |
| *Ccr2*<sup>-/-</sup> CD45.1<sup>+</sup>/CD45.2<sup>+</sup> | The Francis Crick Institute | (Kuziel et al., 1997) |

**Table 2.**
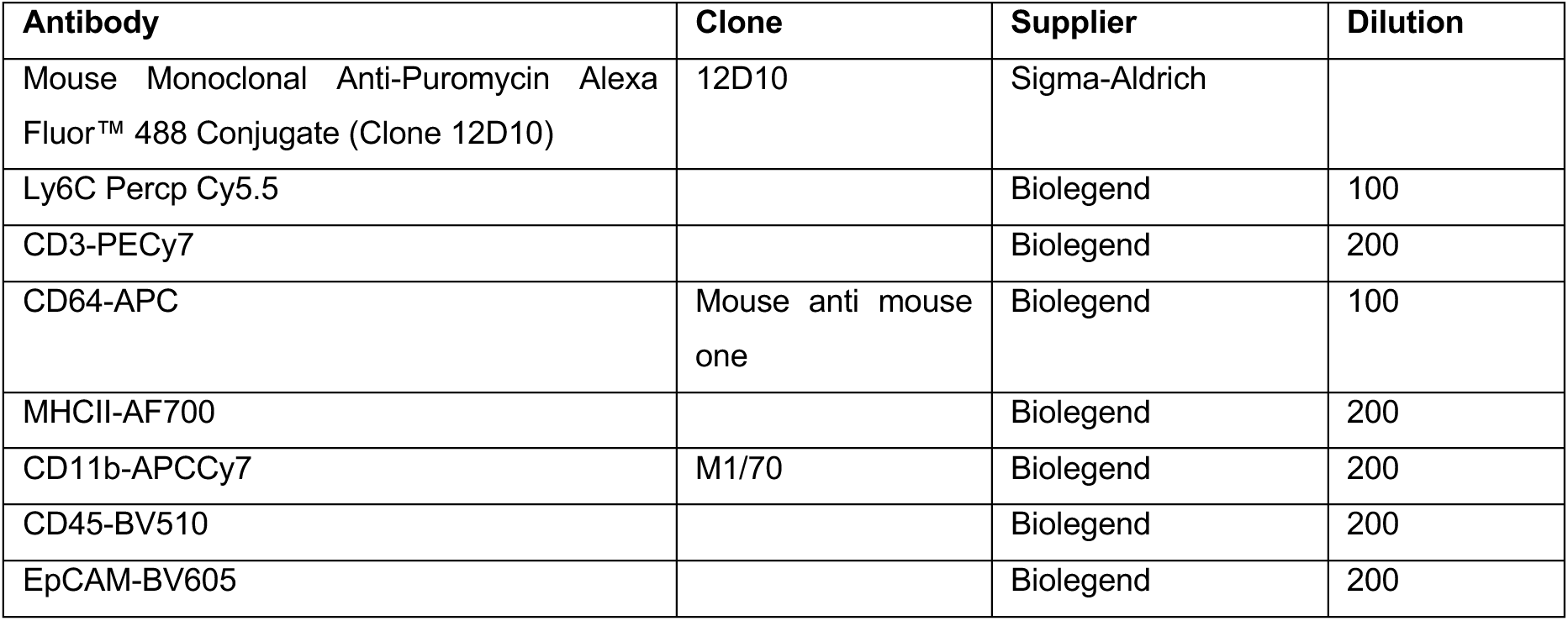
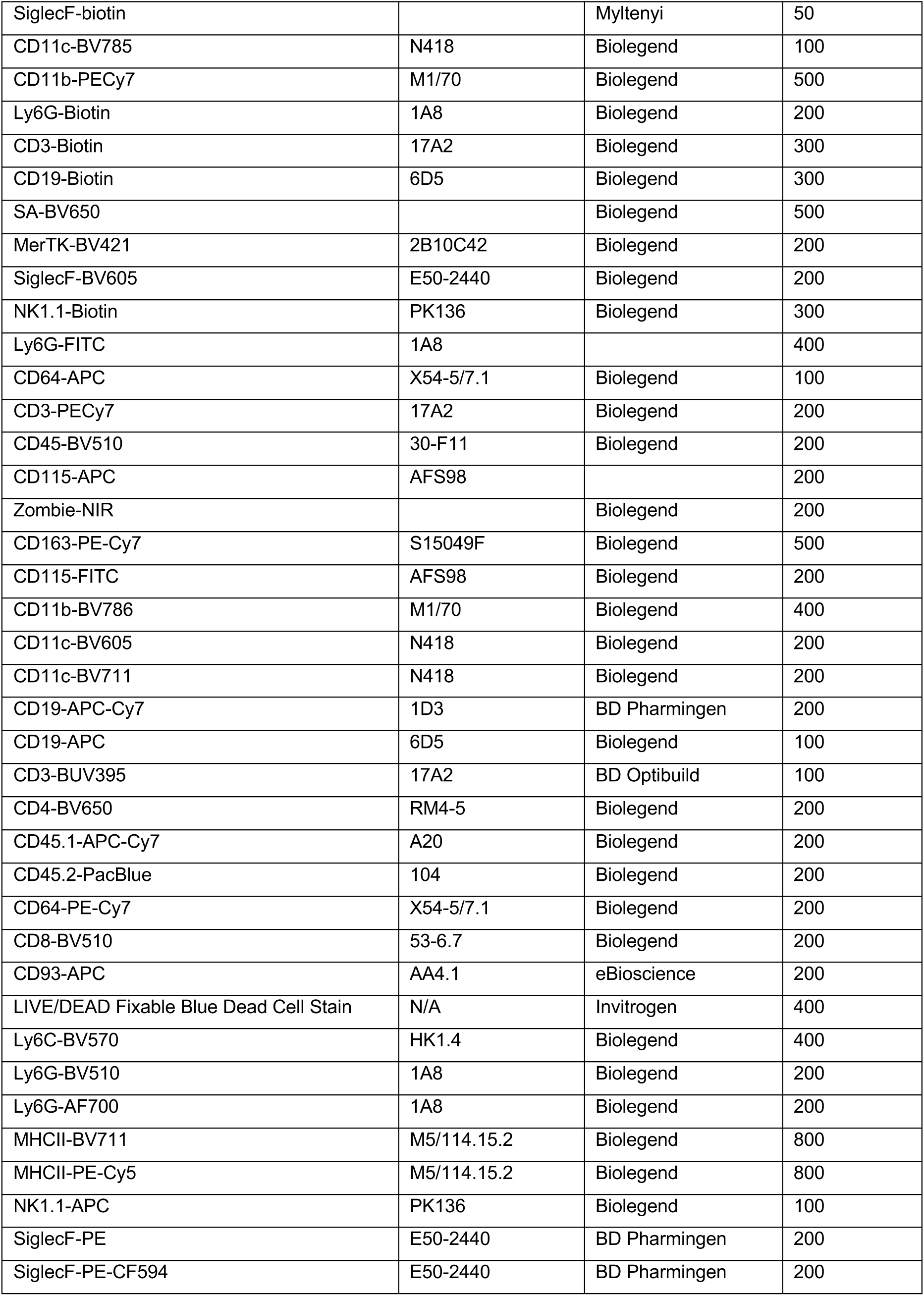
Flow cytometry and Fluorescence Activated Cell Sorting antibodies.

### RSV stock generation and plaque assay

RSV strain A2 (kindly provided by Dr James Harker; Imperial College) was expanded in HEp-2 cells (ECACC - CVCL_1906) in RPMI containing 100 U/ml Pen/Strep (Gibco), 2 mM L-Glutamine (Gibco). When syncytia were observed throughout the culture and the confluency dropped by 10-20 %, cells were harvested, sonicated and vortexed to promote release of viral particles. Following centrifugation at 900 g for 10 minutes (4 °C), supernatant was aliquoted and stored in a liquid nitrogen tank. RSV titre was assessed by immuno-plaque assay as previously described^59^. To generate UV-inactivated RSV (UV-RSV), virus suspensions were transferred to a 48-well plate and irradiated with a UV-C dose of 2 J/cm^2^ using a Spectrolinker XL-1500 UV Crosslinker (Spectro-UV).

### Infections

Mice were infected with Influenza A Virus X31 (10000 TCID_50_ in 30 μl PBS) or RSV A2 (5 x 10^5^PFU in 50 μl RPMI via intranasal instillation under light anaesthesia (3% isoflurane). Subsequent challenge with *S. pneumoniae* (strain TIGR4) was performed by inoculating mice with 5 x 10^5^ CFU diluted in RPMI in 30μl intranasally. Preinfection body weights were recorded and mice were weighed daily (at similar times of day) and monitored for clinical symptoms. Mice reached clinical endpoint at 25% weight loss from baseline weights or upon achieving a clinical score equivalent or exceeding 5 points. Clinical scores were determined by assigning 1 point for each of the following: piloerection, hunched posture, labored breathing, decreased movement, movement only on provocation, partially closed eyes, or hypothermia. No movement on provocation or middle ear infection (disrupted balance) resulted automatically in a score of 5 and mice culled immediately.

### Bone marrow chimeric mice

Bone marrow of recipient mice was depleted by irradiation or busulfan injection. To generate irradiation chimeras, recipient mice (C57BL/6 CD45.2) were anaesthetised using 1 mg/kg Medetomidine Hydrochloride (Dormitor) and 75 mg/kg ketamine (Ketavet) in 0.9% Saline (Thermo Fisher Scientific). Mice were irradiated with 10 Gy with thorax and head protected by a lead shield. After 20 minutes mice received anti-sedan reversal agent and were placed on a heat mat in their cages for 40 minutes. The next day, approximately 3×10^6^ donor bone marrow cells from C57BL/6 CD45.1/2 were transferred into recipient mice by tail vein injection. Successful reconstitution was assessed by blood sampling at 6-8 weeks post bone marrow transfer.

Alternatively, bone marrow was depleted by i.p. injection of two doses of 10 mg/kg busulfan (Busilvex, Pierre Fabre) with 24 hours between injections. Around 24 hours after the last injection, 10-20×10^6^ donor bone marrow cells were administered via tail vein injection. Successful reconstitution was assessed by blood sampling at 6-8 weeks post bone marrow transfer.

### EdU incorporation

Mice were injected intravenously with 1mg EdU and BALF collected 24 hours later. Cells were subjected to surface staining followed by treatment with the Click-iT EdU Cell Proliferation Kit for Imaging, AlexaFluor 647 dye (Invitrogen).

### Lung processing for flow cytometry

Mice were euthanised by i.p. overdose of Dolethal (Vetoquinol). Blood was flushed from the lungs by injecting 10 ml PBS through the right cardiac ventricle. Lungs were excised and placed in ice-cold RPMI1640 + 10 % FBS until further processing. Lungs were placed in a 7 ml bijou container and roughly cut with scissors. Next, 2 ml pre-warmed RPMI1640 + 10 % FBS supplemented with 0.4 mg/ml collagenase V (Sigma), 0.625 mg/ml collagenase D (Roche), 1 mg/ml dispase (Gibco) and 30 μg/ml DNase (Roche). Lungs were digested for 25 minutes at 37 °C in a shaking incubator at 200 rpm. Digested tissue was passed through a 100 μm filter and centrifuged at 400 g for 5 minutes (4 °C). Red blood cells (RBC) were lysed by resuspending the cell pellet in 3 ml RBC lysing buffer (Sigma) and incubating for 3 minutes at room temperature. RBC lysis was stopped by addition of 15 ml FACS buffer and samples were centrifuged at 400 g for 5 minutes (4 °C). Finally, pellets were resuspended in 1 ml FACS buffer (PBS + 2 mM EDTA + 2 % FBS) and passed through a 40 μm filter.

### Broncho-alveolar lavage processing of flow cytometry

Lungs were inflated 3-7 times with either ice-cold or pre-warmed PBS (37 °C) + 2 mM EDTA (Thermo Fisher Scientific). The first BAL was centrifuged at 400 g for 5 minutes (4 °C) and supernatant was stored at -80 °C. The pellet of the first BAL was pooled with subsequent washes, centrifuged at 400 g for 5 minutes (4 °C) and resuspended in FACS buffer.

### Blood processing for flow cytometry

Between 100-500 μl of blood was taken from the inferior vena cava and mixed with 20 μl 0.5M EDTA (Gibco) in a 15 ml Falcon tube placed on ice. RBC lysis was performed by adding 1 ml 1x RBC lysis buffer (Biolegend) to the blood, vortexing vigorously and incubating for 5 minutes on ice. After 5 minutes, 10 ml of FACS buffer was added and cells were pelleted by centrifugation at 400 g for 5 minutes (4 °C). Supernatant was removed and RBC lysis was repeat two more times. After the final lysis, the pellet was resuspended in FACS buffer.

### Histology

Sample preparation for histology: Mice were euthanized with 600mg/kg of pentobarbital and were subsequently perfused via the right ventricle with 10mL of PBS. Lungs were inflated via the trachea with 1mL 10% neutral buffered formalin (NBF), and 5 lobes were dissected and placed in a bijoux containing 5mL 10% NBF and fixed overnight at room temperature. Lungs were then transferred to 70% ethanol before embedding in paraffin wax and sectioning. Sections were stained with haematoxylin and eosin (H&E).

Histopathology analysis: Scoring was performed, blinded to group, by board-certified veterinary pathologists Simon Priestnall and Alejandro Suarez-Bonnet from the Royal Veterinary College/Francis Crick Institute. A semi-quantitative scoring method was applied for histopathological observations: 0 = no lesion; 1 = minimal; 2 = mild; 3 = moderate, 4 = marked changes. In addition, observations were normalized by multiplying the proportion of tissue affected by the score for a given observation. Histopathological findings were defined as follows: vascular leakage = intra-alveolar protein (including fibrin) or oedema; epithelial damage = erosion, ulceration, abscess; pleural thickening = reactivity/hypertrophy; perivascular cellular infiltration = lymphocytes/plasma cells, polymorphonuclear cells; alveolar damage and cellular infiltration = thickening of septae (including type II pneumocyte hyperplasia), AMs and neutrophils.

### Bone marrow processing for flow cytometry

Femur and tibia of both legs were collected in Air-Buffered Iscove’s Modified Dulbecco’s Medium (AB-IMDM, prepared in-house). Muscle and tendons were removed aseptically, and incisions made on proximal and distal ends of each bone. To allow flow-through of BM cells, a 21-gauge needle was used to generate a hole at the bottom center of the 0.5 mL tube. Bones were placed together with 0.3 mL of AB-IMDM into the punctured 0.5 mL collection tube, which was placed into 2 mL collection tube. Cells were pulse-centrifuged out of the bone at 10,000 rpm on a benchtop centrifuge, and were passed through a 70μM strainer. Red blood cells were lysed with 5mL 0.83% ammonium chloride at RT for one minute, and the reaction halted with 25 mL of AB-IMDM. Cells were washed and counted in 45 mL PBS, and then spun down and resuspended to the desired volume in PBS for I.V. injection.

### *In vitro* challenge

Where bulk AMs were restimulated, AMs were counted based on morphology within whole bronchoalveolar lavage fluid using a hemocytometer and cells were plated at a density of 50,000 AMs per well. AMs were allowed to adhere for one hour, followed by two washes with warm PBS to remove non-adherent cells. The remaining cells represented AMs with >95% purity. Where fAMs and mono-AMs were separated, cells were separated as indicated using fluorescence activated cell sorting, and AMs were subsequently plated at 50,000 cells per well. Stimulation was performed where indicated with 100 ng/mL LPS (Enzo), 100ng/mL Pam3CSK4 (Enzo), 1 mM CpG (Invivogen tlrl-2395), 1 mg/mL 3p-hpRNA (Invivogen tlrl-hprna) complexed with Lyovec (Invivogen) according to manufacturer’s instructions, or poly(I:C) (Invivogen tlrl-pic) in a total volume of 200ml. Supernatants were collected after 16h and cells lysed in 200 ml RLT lysis buffer (Qiagen) with b-Mercaptoethanol (Sigma) and stored at 80 °C until RNA extraction.

### Luminex assay

Supernatant from first wash of bronchoalveolar lavage or supernatants from cell culture were used for 26-plex cytokine and chemokine array using ProcartaPlex Mouse Cytokine & Chemokine Panel 1 (Luminex - EPX260-26088-901), following manufacturer’s instructions. IFN-α and -β were quantified using ProcartaPlex Mouse IFN-α/IFN-β panel, 2plex (Invitrogen - EPX02A-22187-901)

### ELISA

Supernatants from *ex-vivo* stimulus were used for ELISA for IL-6 (Invitrogen – 88-7064-88) and TNF (Invitrogen – 88-7324-88) following manufacturer’s instructions. Supernatants from bulk AMs stimulated with LPS, Pam3CSK4 and CpG were diluted 1:6 before ELISA.

### Intracellular cytokine staining

For intracellular cytokine staining, cells were washed with PBS after fixation with PFA as described above and resuspended in permeabilization buffer (eBioscience, 10x permeabilization buffer from Foxp3 TF staining buffer set – 00-5523-00) for 25 minutes at RT in the dark. Cells were then stained with anti-cytokine antibodies for 1 hour in permeabilization buffer + Fc-Block at RT in the dark. Cells were washed in PBS and resuspended in PBS for flow cytometry analysis.

### SCENITH

AMs in bronchoalveolar lavage fluid were counted and total cells were plated at a density of 100,000 cells per well. After one-hour, non-adherent cells were removed by washing twice with warm PBS and cells were resuspended in RPMI +/- 100ng/mL Pam3CSK4 (Enzo). After two hours, SCENITH protocol was initiated, adapted from (REF). Cells were either treated with 2M 2-Deoxy-D-Glucose (Merck - D6134-25G), 1mM Oligomycin (Merck - 75351-5MG), a combination of 2-Doexy-D-Glucose and oligomycin, 5mg/mL Harringtonin (AbCam AB141941), or no inhibitor, before application of 10mg/mL puromycin (Merck - P7255). 2-Deoxy-D-Glucose and Harringtonin were applied 15 minutes before puromycin, while oligomycin was applied 5 minutes before puromycin. 35 minutes after application of puromycin, cells were washed with PBS and incubated with accutase (Stempro) for 15 minutes. Cells were detached by pipetting and transferred to a V-bottom 96-well plate for surface staining, followed by intracellular/intranuclear staining with a-Puromycin-AF488 (Clone 12D10) using the eBioscience Foxp3 / Transcription Factor Staining Buffer Set following manufacturer’s instructions.

### Flow cytometric metabolic assays

In-vitro metabolic assays were performed as previously described (Crotta et al., 2023). Briefly, uptake of fatty acids and glucose was measured by exposure of cells to 0.5μM Bodipy-C16 (4,4-Difluoro-5,7-Dimethyl-4-Bora-3a,4a-Diaza-s-Indacene-3-Hexadecanoic Acid) (Invitrogen – D3821) or to 20 μM 2-NBDG (Invitrogen – N13195) for 30 min at 37 °C, 5 % CO_2_. Labelling of mitochondria was done by exposure of cells to cell permeant Mitotracker Orange (Molecular Probes – M7510) (100 nM) or to TMRM (Invitrogen – T668) (50nM) for 30 minutes at 37 °C, 5 % CO_2_. For all assays, cells were washed three times with PBS + 2 % FCS, followed by surface staining and analysis by flow cytometry without fixation.

### Clodr-lipo

50μl clodr-lipo or control PBS liposomes (Liposoma) were administered intratracheally under light (isofluorane-induced) anaesthesia twice at a 24-hour interval.

### Gene expression analysis by qPCR

Lung tissue was stored in RNA*later* (Invitrogen) on ice for 2-6 hours, transferred to a PCR-clean 2 ml Eppendorf tube and stored at -80°C until further processing. Before starting the RNA extraction, 650 μl of RLT Buffer Plus (QIAGEN) containing 1% β-mercaptoethanol (Sigma) and a 0.4 cm stainless steel bead (QIAGEN) was added to each tube. Samples were homogenised (2 x 1 min @ 25/s) with a TissueLyser (QIAGEN). RNA was extracted using the RNeasy micro kit plus (QIAGEN) following manufacturer’s instructions. RNA (1μg) was converted into cDNA using the High-capacity cDNA reverse transcription kit (Applied Biosystems) following manufacturer’s instructions. Gene expression was assessed by qPCR using the Sensifast SYBR lo-rox kit (Bioline) with primers (Integrated DNA Technologies) listed in **Table 3**. Amplification and quantification were performed using the Quantstudio 5 (Thermo Fisher Scientific). Gene expression was normalised to *B2m* and relative gene expression was calculated using the 2^-ΔΔC(t)^ method.

**Table 3.**
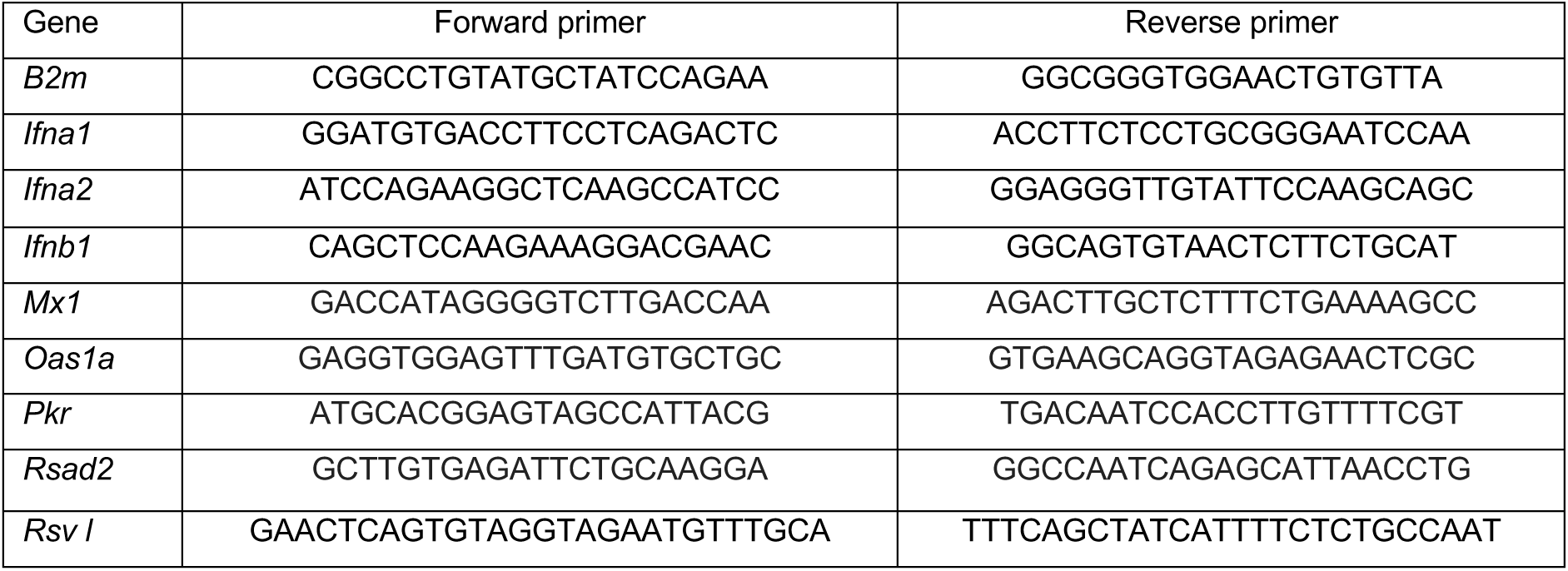
qPCR primers.

### Sample preparation for bulk RNA sequencing

RNA sequencing data included was performed in three separate experiments and sequencing runs. The first included 1mo and 2mo RSV timepoints, the second the 4mo RSV timepoint, and the third 12d, 1mo clodronate and 1mo IAV timepoints. RNA was extracted from frozen lysates of overnight stimulated sorted AMs using the RNeasy Micro Kit (Qiagen - 74004) according to manufacturer’s instructions, with on-column DNase digestion using RNase-free DNase (Qiagen - 79254). RNA integrity was confirmed by RNA electrophoresis with TapeStation system (Agilent). Libraries were prepared using NEBNext Low Input Ultra II RNA Library Prep Kit (Illumina). Sequencing was carried out on the Illumina NovaSeq6000 and typically generated ±25 million 100bp paired-end reads per sample.

### Bulk RNA sequencing bioinformatics analysis

Library preparation: Bioinformatic analysis of fastq files was performed using the nf-core/rnaseq pipeline version 3.8.1 in Nextflow V21.10.3 with default parameters. Briefly, re-sequenced FastQ files were merged (cat), QC performed (FastQC), followed by adapter and quality trimming (Trim Galore!). Reads were then aligned (STAR) to the mouse genome version mm10 (mouse Ensembl GRCm38 - release 95) and quantified (rsem).

Analyses were performed with R (v4.3.0) and ggplot2 v3.4.2 was used for data visualisation. Differential gene expression: Differential gene expression analysis was performed using DESeq2 (v1.42.0). Ensembl gene identifiers were mapped to gene symbols using the org.Mm.eg.db (v3.17.0) annotation package. Lowly expressed genes were filtered out prior to testing, retaining only genes with at least 10 counts in a minimum of two samples. Normalization and dispersion estimation were performed using DESeq2’s default median-of-ratios method and shrinkage estimators. Wald tests were used to assess differential expression between conditions. P-values were adjusted for multiple testing using the Benjamini–Hochberg procedure to control the false discovery rate (FDR). Genes were considered significantly differentially expressed if they met an adjusted p-value (FDR) threshold of < 0.05 and an absolute log2 fold-change > 1.

Principal component analysis (PCA): Top 500 most variable genes were input and genes with zero variance across samples were removed. PCAwas performed on stimulated and mock treated samples separately, using the prcomp function in base R with centring and scaling enabled.

Clustering and heatmap: Gene expression data for the top 500 differentially expressed genes (no-stimulation condition) were imported. Rows containing missing values and genes with low expression variability (lowest 25% variance) were excluded. Pairwise sample distances were calculated using the Spearman correlation distance metric (amap package, v0.8-19), and hierarchical clustering was performed using average linkage (hclust). An initial clustering with *k* = 10 groups was used to identify and remove clusters containing fewer than 15 genes. The remaining genes were re-clustered, and the final dendrogram was cut into four clusters (*k* = 4). The filtered and ordered expression matrix was visualized using the pheatmap package (v1.0.12) with row and column ordering based on cluster membership and treatment group, respectively, and colour representing row-wise z-score.

Pathway analysis: Each gene cluster was intersected with differentially expressed genes at one- and two-months post-RSV. Over-representation analysis (ORA) was performed on the resulting gene lists using clusterProfiler (v4.8.1) and org.Mm.eg.db (v3.17.0) for Gene Ontology (GO) Biological Process terms. Enrichment *p*-values were adjusted for multiple testing using the Benjamini–Hochberg method. Enrichment results were simplified to remove redundant GO terms, retaining the most significant term per cluster. The top three significantly enriched terms per cluster (adjusted *p* < 0.05) were visualized showing −log10(adjusted *p*-value) scores.

Data normalization and integration across experiments: To enable direct comparison of transcriptional responses across independent experiments, datasets were merged and normalized. First, gene identifiers were harmonized by matching Ensembl IDs across datasets, retaining only genes common to all experiments. To account for baseline differences across experiments, counts were normalized relative to corresponding PBS-treated control samples within each batch. To further correct for subtle inter-batch effects associated with stimulation differences, a batch correction factor was derived by comparing PBS-treated mock and stimulated samples within each experiment and this ratio was used to scale stimulated samples.

### Single cell RNA sequencing analysis

Data from Li et al. 2022^35^ describing monocyte/macrophage diversity and dynamics following influenza were downloaded and underwent standard processing. Low quality cells removed using mitochondrial reads and low feature number. Doublets removed using DoubletFinder. Data were then integrated with Harmony. Cell types labelled using canonical features and superclusters defined by manual annotation using signature from McCowan et al. To perform module scoring, gene signatures were generated from our bulk RNA-sequencing and enrichment assessed across superclusters.

### Tamoxifen administration

Labelling of CCR2^+^ cells was achieved by administering one dose of 5 mg tamoxifen (Sigma Aldrich) in 100 μl sesame oil (Sigma) or corn oil (Sigma) by oral gavage.

### Statistical analysis

Statistical analysis was performed using GraphPad Prism 10 (GraphPad). Statistical tests are stated in Fig. legends and were selected based on number of experimental groups, data distribution and variance characteristics. The investigators were not blinded to the mouse treatment group allocation.

## List of instruments

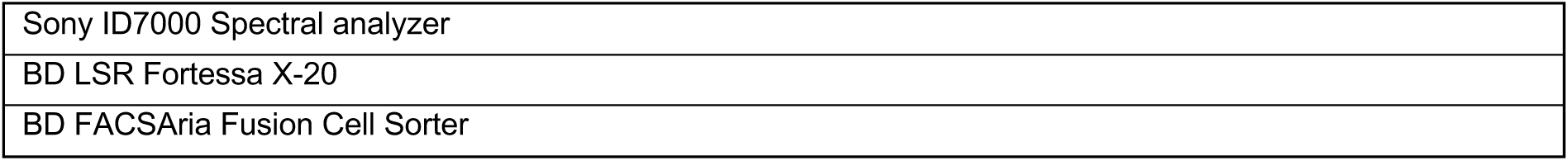

|  |
| --- |
| Sony ID7000 Spectral analyzer |
| BD LSR Fortessa X-20 |
| BD FACSAria Fusion Cell Sorter |

## List of reagents, chemicals

| Reagent | Supplier | Cat # |
| --- | --- | --- |
| 123count eBeads™ Counting Beads | Invitrogen | 01-1234-42 |
| 2-Deoxy-D-Glucose | Merck | D6134-250MG |
| 2-NBDG (2-(N-(7-Nitrobenz-2-oxa-1,3-diazol-4-yl)Amino)-2-Deoxyglucose) | Invitrogen | N13195 |
| 3p-hpRNA | Invivogen | tlrl-hprna |
| Accutase | Stempro | A1110501 |
| Iscove's Modified Dulbecco's Medium | Gibco | 12440053 |
| Ammonium Chloride | Sigma | 213330 |
| BODIPY™ FL C16 (4,4-Difluoro-5,7-Dimethyl-4-Bora-3a,4a-Diaza-s-Indacene-3-Hexadecanoic Acid) | Invitrogen | D3821 |
| Busulfan | Pierre Fabre |  |
| Clodr-lipo | Liposoma | CP-005-005 |
| Collagenase D from Clostridium histolyticum | Roche | 11088858001 |
| Collagenase from Clostridium histolyticum, Type V | Sigma-Aldrich | C9263 |
| Corn oil | Sigma | C8267 |
| CpG | Invivogen | tlrl-2395 |
| Dispase II | Gibco | 17105041 |
| DNase I grade II, from bovine pancreas | Roche | 10104159001 |
| eBioscience™ Foxp3 / Transcription Factor Staining Buffer Set | Invitrogen | 00-5523-00 |
| Ethanol | Sigma-Aldrich | 51976 |
| Foetal Bovine Serum |  |  |
| GM-CSF |  |  |
| Harringtonine | AbCam | ab141941 |
| L-glutamine | Gibco | 25030081 |
| LPS | Enzo |  |
| Lyovec | Invivogen | lyec-1 |
| MitoTracker Orange CMTMRos | Invitrogen | M7510 |
| Mouse IL-6 Uncoated ELISA Kit | Invitrogen | 88-7064-88 |
| Mouse TNF alpha Uncoated ELISA Kit | Invitrogen | 88-7324-88 |
| NEBNext® Ultra™ II RNA Library Prep Kit for Illumina® | New England Biolabs | E7770L |
| 10% Neutral Buffered Formalin |  |  |
| Oligomycin | Merck | 495455 |
| PAM3CSK4 | Enzo | ALX-165-066 |
| PBS liposomes | Liposoma | CP-005-005 |
| PBS | Gibco | 10010023 |
| Penicillin-Streptomycin | Gibco | 15140122 |
| Poly(I:C) | Invivogen | tlrl-pic |
| ProcartaPlex Mouse Cytokine & Chemokine Panel 1 | Luminex | EPX260-26088-901 |
| Puromycin | Merck | P7255 |
| RBC Lysis Buffer 10x | Biolegend | 420302 |
| RNAlater™ Stabilization Solution, 500 mL | Invitrogen | AM7021 |
| RNase-Free DNase Set | Qiagen | 79254 |
| RNeasy Micro Kit | Qiagen | 74104 |
| RNeasy Plus Micro Kit | QIAGEN | 74034 |
| RNeasy Plus Mini Kit | Qiagen | 74136 |
| RPMI 1640 | Gibco | 11875093 |
| Sensifast Sybr-lo ROX master mix | Bioline | BIO-94020 |
| Sesame oil | Sigma | S3547 |
| Tamoxifen | Sigma-Aldrich | T5648-1G |
| Tetramethylrhodamine, Methyl Ester, Perchlorate (TMRM) | Invitrogen | T668 |
| Transforming Growth Factor-β | Peprtech | 100-21 |
| UltraComp eBeads™ Plus Compensation Beads | Invitrogen | 01-3333-42 |
| UltraPure™ 0.5M EDTA, pH 8.0 | Invitrogen | 15575020 |
| β-mercaptoethanol | Sigma-Aldrich | M3148 |

## Supporting information

Combined Supplementary Data

## Acknowledgements

The authors are grateful to Dr. James Harker (Imperial College, London) for provision of RSV stocks. Thanks are paid to Cameron Bullen for invaluable bioinformatics support. We are also grateful for fruitful discussions with Dr. Will Branchett and Prof. Anne O’Garra about this work. We are grateful to Keshav Sharma for methodological support with EdU and Annexin V staining. The authors gratefully acknowledge the IRR Flow Cytometry and Cell Sorting Facility (University of Edinburgh), MVLS-SRF Flow Cytometry Core Lab (University of Glasgow), the Crick Flow Cytometry Science and Technology Platform (The Francis Crick Institute), and the Crick Experimental Histopathology Science and Technology Platform for their support and assistance in this work. We are grateful to the Genomics Science and Technology Platform at the Crick for performing library preparation and RNA sequencing and to Dr. Gavin Kelly and Dr. Miriam Llorian-Sopena at The Francis Crick Institute for RNA-sequencing data processing and analysis. We would like to thank the Bioresearch and Veterinary Services at the University of Edinburgh, Biological Services at the University of Glasgow and the Biological Research Facility at The Francis Crick Institute for husbandry of our mice and other technical assistance. Servier Medical Art was used for the generation of some of the graphics. **This research was funded in part by the Wellcome Trust**. **For the purpose of open access, the author has applied a CC BY public copyright licence to any Author Accepted Manuscript version arising from this submission.**

## CRediT Author Contributions

CRedit Author Contributions.
|  | Conceptualisation | Data Curation | Formal Analysis | Funding Acquisition | Investigation | Methodology | Project Administration | Resources | Software | Supervision | Validation | Visualization | Writing – original draft | Writing –review & editing |
| --- | --- | --- | --- | --- | --- | --- | --- | --- | --- | --- | --- | --- | --- | --- |
| CSI | X | X | X |  | X | X |  |  |  |  | X | X | X | X |
| WT'J | X | X | X | X | X | X |  |  |  |  | X | X | X | X |
| ICM |  | X | X |  | X | X |  |  |  |  | X | X | X | X |
| SB |  |  | X |  |  |  |  |  |  |  |  |  |  | X |
| JL |  |  | X |  | X |  |  |  |  |  |  |  |  | X |
| G-RJ |  |  |  | X | X | X |  |  |  |  |  |  |  | X |
| JK |  |  |  |  | X | X |  |  |  |  |  |  |  | X |
| SC |  |  |  |  | X | X |  |  |  |  |  |  |  | X |
| SP |  |  | X |  | X |  |  |  |  |  |  |  |  | X |
| AS-B |  |  | X |  | X |  |  |  |  |  |  |  |  | X |
| JS | X |  |  | X |  | X |  |  |  |  |  |  |  | X |
| AW | X |  | X | X |  | X | X | X |  | X | X | X | X | X |
| CCB | X | X | X | X | X | X | X | X |  | X | X | X | X | X |

## Data and material availability

All data needed to evaluate conclusions in the paper are present in the paper or Extended Data Materials, and the RNA-seq data will be desposited in the National Center for Biotechnology Information Gene Expression Omnibus public database (www.ncbi.nlm.nih.gov.geo/) upon publication.

