## Supplementary material for "Diverse lung challenges elicit a conserved monocyte-to-macrophage differentiation blueprint": Combined Supplementary Data

### Extended Data Fig. 1

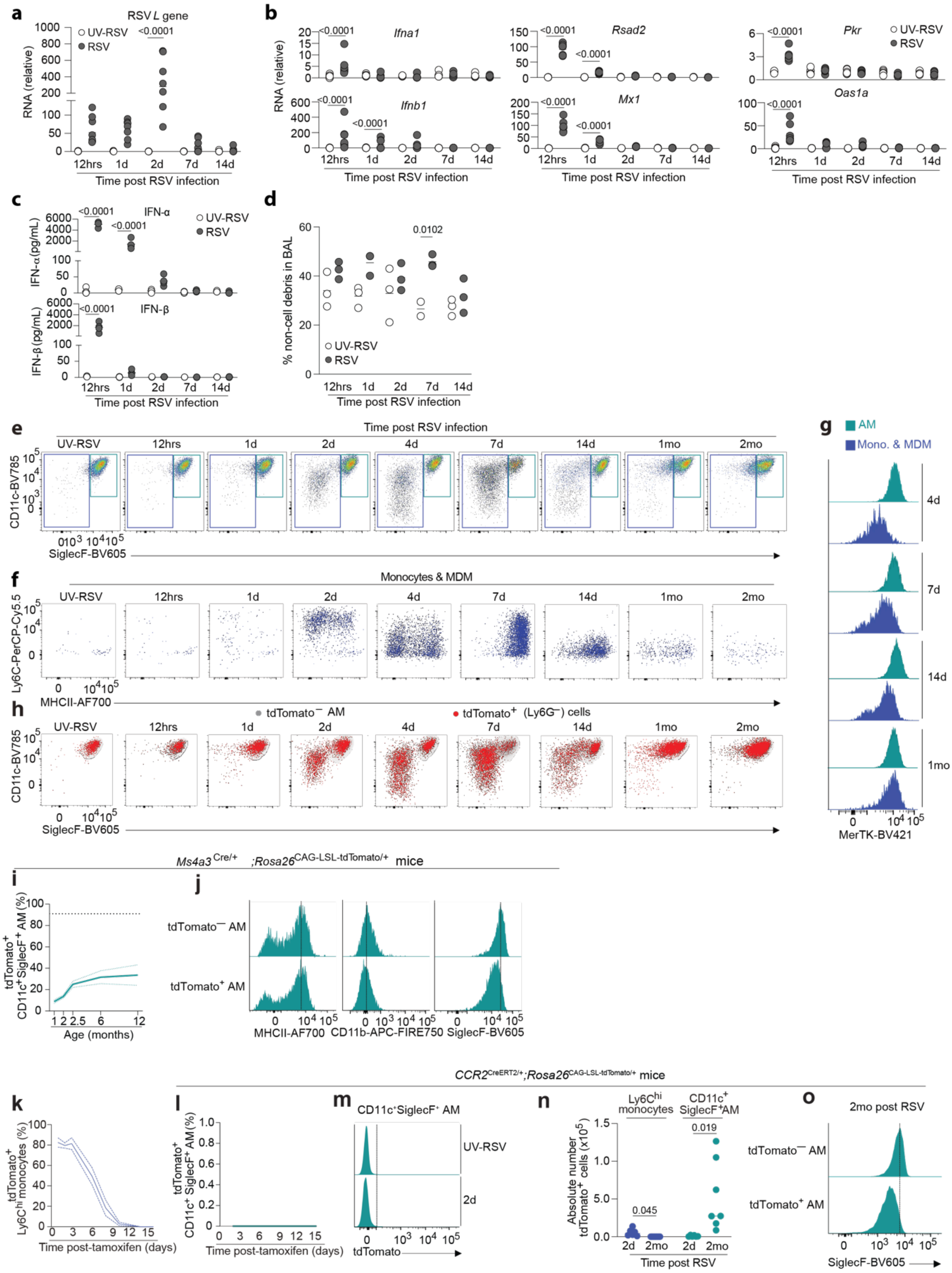


**Extended Data Fig. 1| Establishing infection kinetics and myeloid cell dynamics after RSV infection**

**a.** Relative quantification of *Rsv-l* RNA levels by RT-qPCR in whole lung tissue obtained from C57BL/6J mice inoculated with 5x10$^5$ PFU RSV or UV-inactivated RSV (UV-RSV). Data shown as relative expression to UV-

RSV at 12 hrs post-infection. Symbols represent individual mice. Data pooled from two independent experiments with *n* = 7 mice per group. Statistical significance calculated by two-way ANOVA.

**b.** Gene expression levels of type I IFN (*Ifna1* and *Ifnb1)* and IFN-stimulated genes (*Mx1, Rsad2, Pkr* and *Oas1a)* determined by RT-qPCR using the same whole lung samples as shown in **a** relative to UV-RSV at 12 hrs post-inoculation. Significance assessed by two-way ANOVA.

**c.** IFN-α and -β concentrations in the bronchoalveolar lavage fluid (BALF) from mice, treated as described in Fig. 1a-b were quantified using an IFN-α/β duplex assay. Data from one experiment shown with *n* = 3-4 per timepoint post-RSV or UV-RSV inoculation.

**d.** Frequency of cellular debris in the BALF from mice harvested at the indicated time points (days post infection; dpi) after RSV or UV-RSV inoculation. Symbols represent individual mice. Data from one experiment shown with *n* = 3 mice per group. Statistical significance calculated using one-way ANOVA.

**e.** Representative flow cytometry plots showing manual gating of monocyte/monocyte-derived macrophage (blue/left gates) and AM (green/right gates). Data pre-gated on MerTK$^{+}$ and/or CD11b$^{+}$ (CD45$^{+}$Ly6G$^{-}$CD3$^{-}$ NK1.1$^{-}$CD19$^{-}$) cells isolated from BALF of *Ms4a3*$^{Cre/+}$;*Rosa26*$^{LSL\text{-}CAG\text{-}tdTomato/+}$ mice at the indicated timepoints post-infection.

**f.** Representative flow cytometry plots showing expression of Ly6C and MHCII by cells in BALF within the monocyte/monocyte-derived macrophage gate shown in (e) at the indicated time points.

**g.** Representative flow cytometry histograms showing expression of MerTK by monocyte/monocyte-derived macrophage and AM in BALF at indicated time points post RSV infection.

**h.** Representative flow cytometry plots showing expression of CD11c and SiglecF by tdTomato$^{-}$ AM and Ly6G$^{-}$ tdTomato$^{+}$ cells within the MerTK$^{+}$ and/or CD11b$^{+}$ compartment.

**i.** Proportion of tdTomato$^{+}$ cells amongst CD11c$^{+}$SiglecF$^{+}$ AM in BALF of unmanipulated Ms4a3$^{Cre/+}$;*Rosa26*$^{CAG\text{-}LSL\text{-}tdTomato/+}$mice at indicated ages. Data represented as average proportion of labelled cells ± S.E.M.. Top dotted line represents average proportion of labelled circulating monocytes in blood (91,04 %). Data from one or two independent experiment with *n* = 2-14 mice per group.

**j.** Representative expression of MHCII, CD11b and SiglecF by tdTomato$^{+}$ and tdTomato$^{-}$ AM subpopulations at 2mo post-infection. Vertical lines represent peak histogram level in tdTomato$^{+}$ AMs.

**k.** Proportion of tdTomato$^{+}$ cells within the classical Ly6C$^{hi}$ blood monocyte population in *Ccr2*$^{Cre\text{-}ERT2/+}$*;Rosa26*$^{CAG\text{-}LSL\text{-}tdTomato/+}$ mice given a single dose of tamoxifen and assessed longitudinally. Solid line represents mean with dotted line indicating ± S.E.M. Data from one experiment with *n* = 4-9 mice per group.

**l.** tdTomato kinetics of CD11c$^{+}$SiglecF$^{+}$ AM from BALF in *Ccr2*$^{Cre\text{-}ERT2/+}$*;Rosa26*$^{CAG\text{-}LSL\text{-}tdTomato/+}$ mice given a single dose of tamoxifen and assessed longitudinally Data represented as average proportion of labelled cells ± 1 S.E.M. Data from one experiment with *n* = 4-5 mice per group.

**m.** Representative histogram of tdTomato expression within CD11c$^{+}$SiglecF$^{+}$ AM in *Ccr2*$^{Cre\text{-}ERT2/+}$*;Rosa26*$^{CAG\text{-}LSL\text{-}tdTomato/+}$ mice given a single dose of tamoxifen 1 day before RSV or UV-RSV inoculation and assessed at 2 days after inoculation.

**n.** Absolute number of tdTomato$^{+}$ cells amongst Ly6C$^{hi}$ monocytes in BALF and CD11c$^{+}$SiglecF$^{+}$ AM in BALF obtained from *Ccr2*$^{Cre\text{-}ERT2/+}$*;Rosa26*$^{CAG\text{-}LSL\text{-}tdTomato/+}$ mice given a single dose of tamoxifen 24 hrs prior to inoculation with UV-RSV or RSV. AMs and Ly6C$^{hi}$ monocytes in BALF were assessed at 2 days and 2 months post-infection. Symbols represent individual mice. Data pooled from one (UV-RSV) or two independent (RSV) experiments with *n* = 2-7 mice total per group.

**o.** Representative histogram of SiglecF expression within CD11c$^{+}$SiglecF$^{+}$ AM within tdTomato$^{-}$ and tdTomato$^{+}$ CD11c$^{+}$SiglecF$^{+}$ AM in *Ccr2*$^{Cre\text{-}ERT2/+}$*;Rosa26*$^{CAG\text{-}LSL\text{-}tdTomato/+}$ mice given a single dose of tamoxifen 1 day before RSV or UV-RSV inoculation and assessed at 2 months after inoculation.

**Extended data Fig. 2**

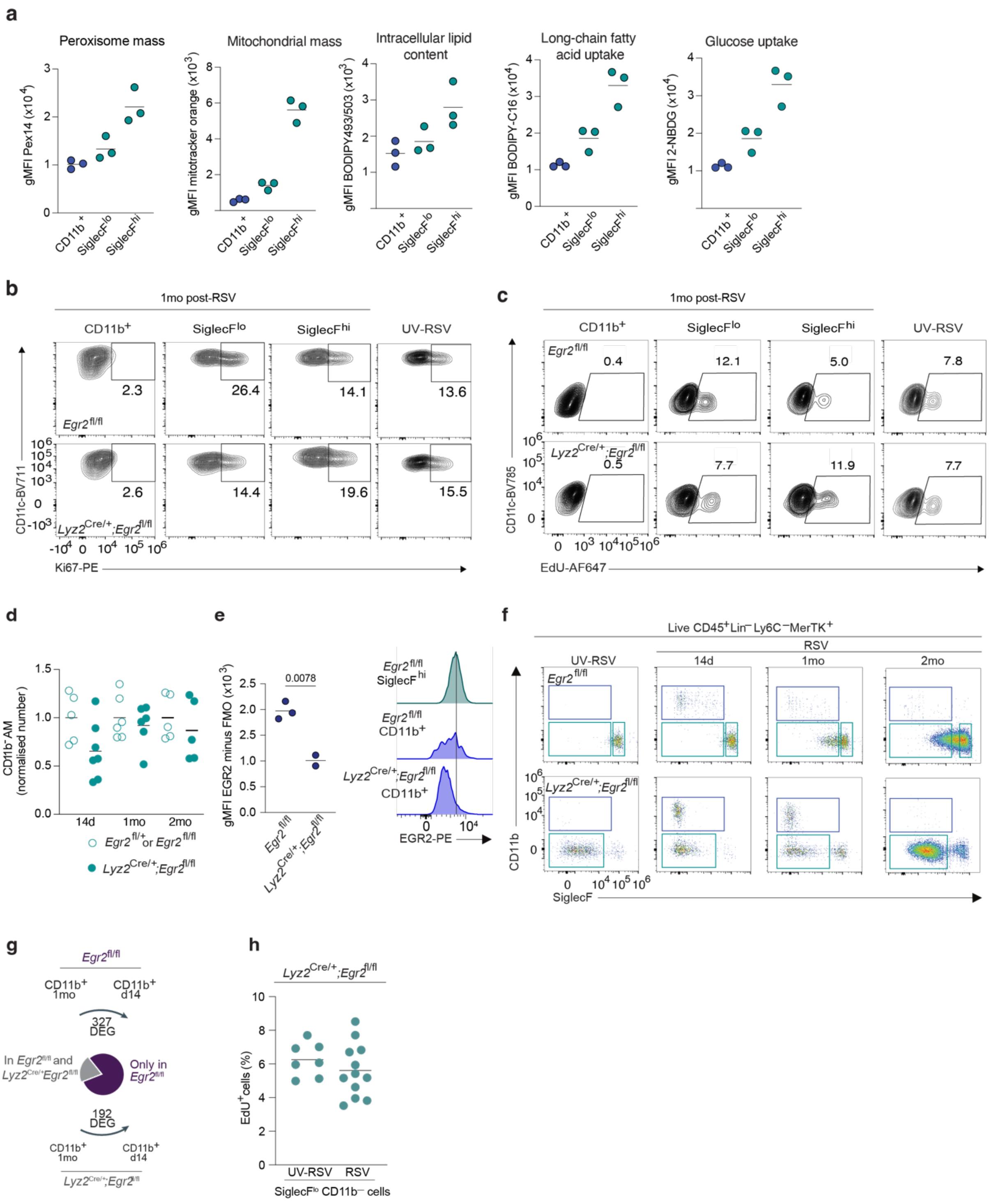


**Extended Data Fig. 2 | Profiling of metabolic activity and EGR2-dependence of AMs**

**a.** Absolute fluorescence values from staining with Pex14-CL594, Mitotracker orange, BODIPY493/503, BODIPY C-16 and 2-NBDG in CD11b- and SiglecF-defined AM subsets. Symbols represent biological replicates. Data representative of two independent experiments with $n$ = 3 per group.

**b.** Representative flow cytometry plots of Ki67 staining in live $CD45^+Lin^-Ly6C^-MerTK^+CD11c^+$ AMs stratified by CD11b- and SiglecF-defined subpopulations from *Egr2*$^{flox/flox}$ control and *Lyz2*$^{Cre/+}$;*Egr2*$^{flox/flox}$ mice infected with RSV.

**c.** Representative flow cytometry plots of EdU incorporation in live $CD45^{+}Lin^{-}Ly6C^{-}MerTK^{+}CD11c^{+}$ AMs stratified by labelled surface marker phenotype from *Egr2*$^{flox/flox}$ control and *Lyz2*$^{Cre/+}$;*Egr2*$^{flox/flox}$ mice infected with RSV. UV-RSV samples derived from fAMs in respective genotypes (SiglecF$^{hi}$ for *Egr2*$^{flox/flox}$ and SiglecF$^{lo}$ for *Lyz2*$^{Cre/+}$;*Egr2*$^{flox/flox}$)

**d.** Normalised number of BALF $CD11c^{+}CD11b^{—}$ AMs from *Egr2*$^{flox/flox}$ control and *Lyz2*$^{Cre/+}$;*Egr2*$^{flox/flox}$ mice. Symbols represent individual mice. Data from *n* = 4-7 mice per group pooled from 3 independent experiments.

**e.** Expression (gMFI) and representative histograms of EGR2 expression by $CD11b^{+}$ cells from *Egr2*$^{flox/flox}$ control and *Lyz2*$^{Cre/+}$;*Egr2*$^{flox/flox}$ mice by flow cytometry. Data from one experiment of *n* = 2-3 mice per group.

**f.** Representative gating strategy used to identify AM subpopulations for RNA-sequencing from *Egr2*$^{flox/flox}$ control and *Lyz2*$^{Cre/+}$;*Egr2*$^{flox/flox}$. Upstream gating on live $CD45^{+}Lin^{-}Ly6C^{-}MerTK^{+}CD11c^{+}$ cells.

**g.** The number of genes differentially expressed in BALF CD11b+ cells between 14d and 1mo post-RSV. The proportion of those differentially expressed *Egr2*$^{flox/flox}$ control which are also differentially expressed in *Lyz2*$^{Cre/+}$;*Egr2*$^{flox/flox}$.

**h.** Frequency of $EdU^{+}$ cells within the SiglecF$^{lo}$ $CD11b^{-}$ compartment after a 24 hour pulse with EdU. *n* = 7-12 mice per group pooled from two independent experiments. Statistically non-significant difference was calculated by a two-tailed T-test.

**Extended Data Fig. 3**

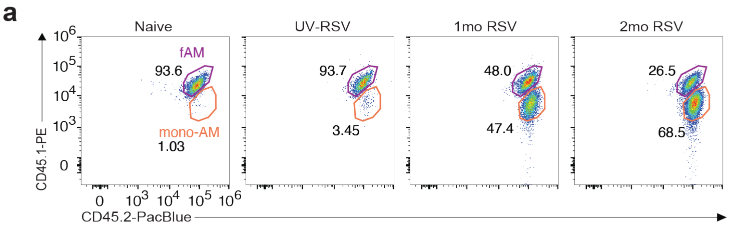


**Extended data Fig. 3 | Identification of mono-AMs in busulfan chimeras after RSV infection.**

a. Representative flow cytometry plots depicting gating strategy used to identify fAMs and mono-AMs within the AM ($MerTK^+CD64^+SiglecF^+CD11c^+$) compartment of busulfan chimeric mice left naive, inoculated with UV-RSV or at one or two months after RSV infection.

### Extended Data Fig. 4

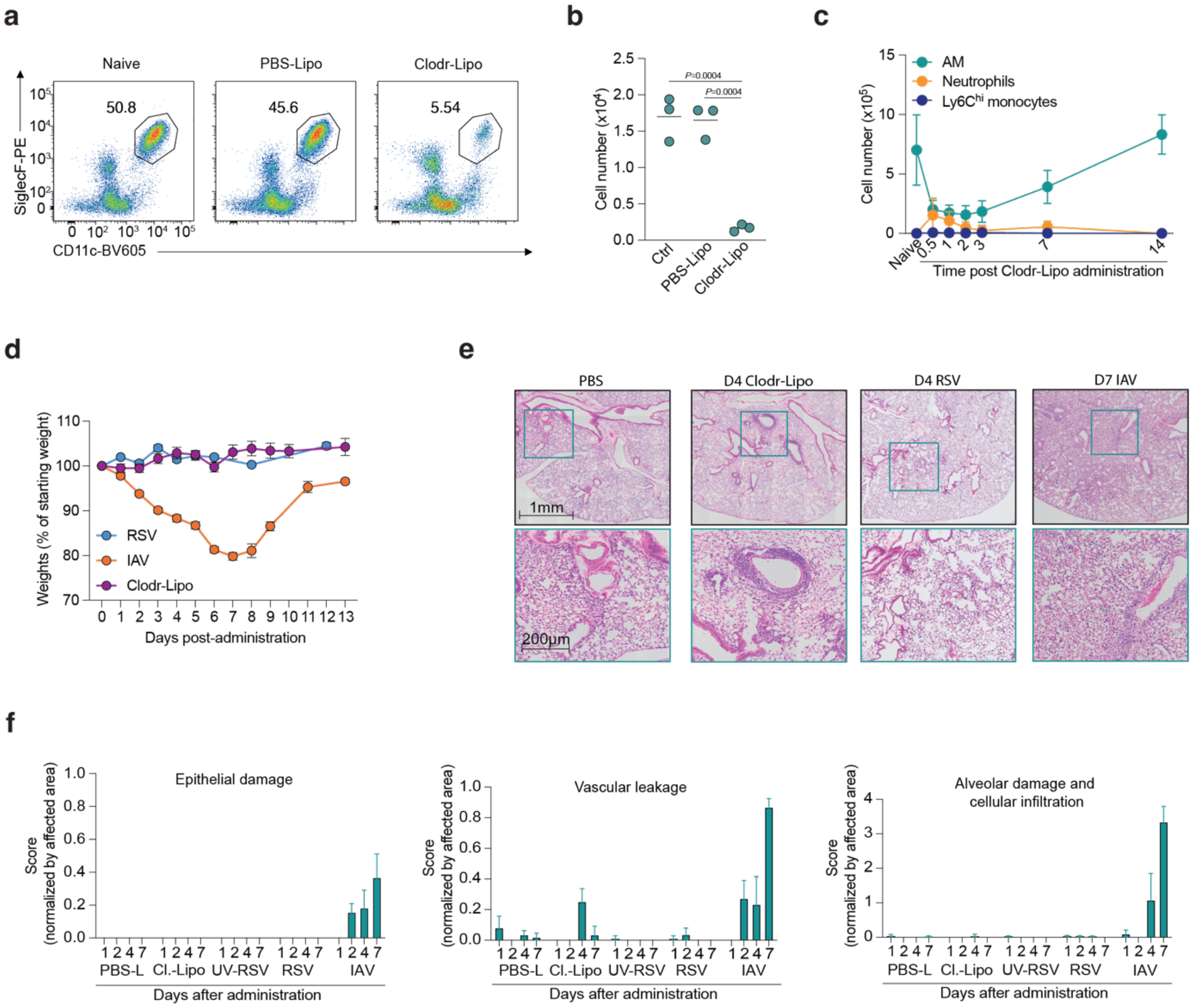


**Extended Data Fig. 4 | Comparative analysis of lung challenge models**

Mice were treated with 50µl clodronate- or PBS liposomes twice intratracheally, or infected with $5 \times 10^5$ PFU RSV (or UV-RSV) or $1 \times 10^4$ $TCID_{50}$ IAV (or PBS).

**a.** Representative flow cytometry plots from whole lungs of mice at 72 hours after clodr-lipo administration showing AM frequency, pre-gated on Live, Ly6G⁻ cells.

**b.** Quantification of AM numbers within whole lungs of mice at 72 hours after clodr-lipo administration. Symbols represent individual mice. Data from one representative experiment of three with $n$ = 3 mice per group. Significance calculated by one-way ANOVA.

**c.** Quantification of AM ($SiglecF^+CD11c^+$, neutrophils ($Ly6G^+$ $CD11b^+$) and $Ly6C^{hi}$ monocyte ($Ly6C^+$ $CD11b^+$) numbers at the indicated timepoints after clodr-lipo administration. Data pooled from 4 independent experiments with $n$ = 3-8 mice per timepoint. Data represented as mean±SD.

**d.** Weight loss of mice infected with RSV, IAV, or treated with clodr-lipo. Symbols represent mean ±SEM from $n$ = 9-22 mice per group per treatment.

**e.** Representative H&E-stained lung section regions (black) with higher magnification segments below (green) from mice administered PBS-lipo, clodr-lipo, UV-RSV, RSV or IAV.

**f.** Histopathological scoring of lung sections stained with H&E from mice administered PBS-lipo, clodr-lipo, UV-RSV, RSV or IAV.. Data represent absolute scores (1-5) multiplied by the proportion (0-1) of the lung affected. Bars represent mean ±SD from n=3-4 mice per group.

### Extended Data Fig. 5

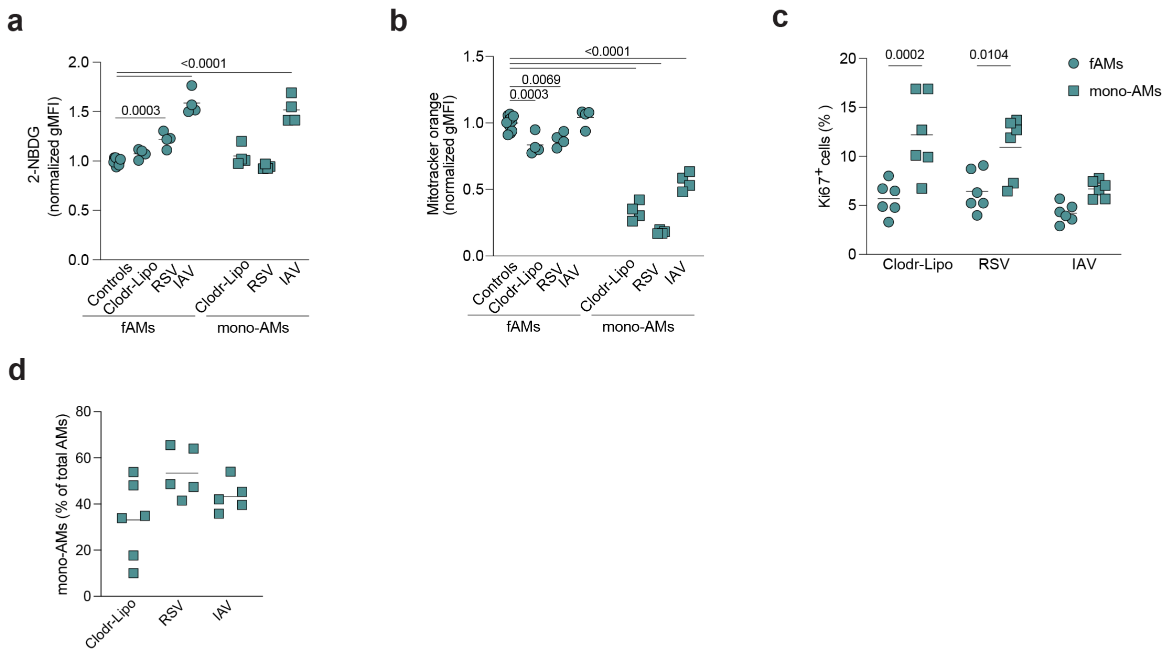


**Extended Data Fig. 5 | Metabolic and proliferative assessment of distinct AM subsets across eliciting contexts**

**a.** Uptake of fluorescent glucose analogue 2-NBDG by AM *in-vitro* from 12d after clodr-lipo administration, 1mo post-RSV and 1mo post-IAV mice, or 12d PBS-lipo, 1mo UV-RSV and naïve controls (pooled together as controls). AM were separated on origin using SiglecF expression. Symbols represent biological replicates. Data pooled from three representative experiments of six, with $n$ = 4 mice per group. Significance assessed by one-way ANOVA.

**b.** Staining with mitotracker orange by AM from 12d after clodr-lipo administration, 1mo post-RSV and 1mo post-IAV mice and 1mo post-IAV mice, or 12d PBS-lipo, 1mo UV-RSV and naïve controls (pooled together as controls). AMs were separated on origin using SiglecF expression. Symbols represent biological replicates. Data pooled from three representative experiments of six, with $n$ = 4 mice per group. Significance assessed by one-way ANOVA comparing all groups to the control condition.

**c.** Ki67 staining of fAM (SiglecF$^{hi}$) and mono-AM (SiglecF$^{lo}$ CD11b$^{-}$) at 12d after clodr-lipo administration, 1mo post-RSV and 1mo post-IAV. Symbols represent biological replicates. Data from one representative experiment of four with $n$ = 6 per group. Significance assessed by two-way ANOVA comparing AM subsets within conditions.

**d.** Frequency of mono-AMs from busulfan chimeric mice at 12d after clodr-lipo administration, or one month after RSV or IAV inoculation. Mono-AMs quantified by frequency of CD45.2$^{+}$ donor-derived cells as a proportion of total CD45.1$^{+}$/CD45.2$^{+}$ cells. Symbols represent biological replicates. Data pooled from two experiments with $n$=5-6 per group. Lack of significance identified by one-way ANOVA.
